# Engineering robust cardiomyocyte differentiation from iPSCs through WNT-mediated process window expansion

**DOI:** 10.64898/2026.09.29.755404

**Authors:** Hirokazu Akiyama, Rin Kobayashi, Shunnosuke Nakamura, Takao Yasui, Kazunori Shimizu, Hiroyuki Honda

**Author notes:** Corresponding author: Hirokazu Akiyama.

## Abstract

Lack of robustness and high variability in cardiomyocyte (CM) differentiation from human induced pluripotent stem cells (hiPSCs) remain critical challenges for cardiac regenerative medicine and biological research. Guided by developmental principles of lineage specification, we investigated whether early WNT inhibition could expand the process window of CM differentiation and thereby improve robustness. To test this, we compared early WNT inhibition from days 1–5 with standard inhibition from days 3–5 following a 1-day CHIR99021 (CHIR) treatment for WNT activation. Focusing on CHIR concentration as a critical parameter, we found that early WNT inhibition substantially broadened the concentration window yielding ≥80% differentiation efficiency across distinct hiPSC lines. Extending the analysis to a two-dimensional parameter space incorporating cell density revealed a markedly expanded ≥80% process window. Temporal analysis of lineage-specific transcription factor expression showed that early WNT inhibition suppressed paraxial mesoderm (PM) specification, likely through *MSGN1* suppression, in the range of high initial WNT activation that otherwise favored PM fate, thereby extending the process window. It also suppressed definitive endoderm (DE) specification within the expanded window, promoting a more favorable lateral plate mesoderm/cardiac fate. We further demonstrated substantially reduced variability under parameter perturbations and across wells and batches, accompanied by higher differentiation efficiency, without compromising CM phenotypes. These findings demonstrate early WNT inhibition as an effective strategy for expanding the process window of CM differentiation and improving robustness, while exemplifying the process engineering principle that a broader window can support more robust differentiation, with potential relevance to stem cell–based bioprocesses.

## 1. Introduction

Cardiovascular disease remains a leading cause of global morbidity and mortality [1]. This substantial disease burden highlights the urgent need for regenerative medicine approaches to overcome the limited regenerative capacity of the adult heart [2,3]. Human induced pluripotent stem cell–derived cardiomyocytes (hiPSC-CMs) represent a promising cell source for cardiac regenerative medicine because they can be generated in virtually unlimited quantities from hiPSCs. Considerable efforts have been made toward the clinical translation of hiPSC-CMs, with several clinical studies underway [2,4,5] and the first hiPSC-CM–based regenerative medicine product receiving marketing approval [6]. Beyond therapeutic applications, hiPSC-CMs have been extensively investigated as human-relevant platforms for disease modeling, drug discovery [7–10], and cardiotoxicity assessment [11–13]. Promoting the broader application of hiPSC-CMs requires continued advances in manufacturing technologies for robust production of high-quality hiPSC-CMs.

For CM differentiation from PSCs, the GiWi protocol pioneered by Lian *et al.* [14,15] has been widely adopted. The protocol is based on biphasic signaling modulation, with a glycogen synthase kinase 3β (GSK3β) inhibitor used for WNT activation followed by a WNT inhibitor. The GiWi protocol offers clear practical advantages because of its simplicity and high efficiency; however, it exhibits substantial well-to-well and batch-to-batch variability in differentiation efficiency, as well as cell line dependence [16–21], all of which complicate the reproducible application of PSC differentiation protocols. Many modifications to CM differentiation protocols have involved incorporating additional cytokines or small molecules, an approach also employed in our prior study, primarily to improve differentiation efficiency [22–26]. However, increasing efficiency through supplementation does not necessarily translate into reduced variability. Meanwhile, to mitigate the impact of differentiation variability on downstream applications, CM enrichment methods have been developed, including post-induction purification based on surface markers [27,28] and metabolic selection [29,30], as well as selection during differentiation [16,31]. Nevertheless, CM enrichment remains a compensatory approach that does not directly address the underlying differentiation instability. Thus, a more fundamental differentiation process design is necessary to enable robust CM manufacturing with both high differentiation efficiency and greater consistency.

In biopharmaceutical and cell therapy manufacturing, understanding the relationships between process parameters and outcomes is a central element of process development [32–34]. The process window can be defined as the range or multidimensional region of parameters over which acceptable performance is maintained. Process robustness refers to the ability to maintain acceptable performance while tolerating variability in process inputs [34]. The window size is one of the key determinants of process robustness, as a broader window provides greater tolerance to sources of variability. This is particularly important for bioprocesses owing to their greater complexity and intrinsic variability compared with those of chemical processes. Among bioprocesses, PSC differentiation is particularly sensitive to various sources of variation, such as the initial cellular state [19], culture materials [35], and culture handling [36], all of which can contribute to differentiation variability. Nevertheless, research in this area is predominantly focused on increasing differentiation efficiency, leaving process robustness largely underexplored. A better understanding and strategic expansion of the process window allowing for efficient differentiation may therefore facilitate the engineering of more robust PSC differentiation processes, with direct applicability to CM differentiation.

In GiWi-based CM differentiation, the WNT-activating GSK3β inhibitor CHIR99021 (CHIR) must be tightly controlled for efficient differentiation because the efficiency is highly dose-dependent and declines outside a narrow concentration range [14,19]. Cell density is another critical parameter that influences cellular states before differentiation [19] and the paracrine environment during differentiation [37]. Both factors can significantly affect differentiation efficiency, potentially by modulating cellular responsiveness to CHIR. The narrowness of the process window defined by these parameters is likely attributable to the critical role of WNT signaling in regulating multiple cell fate decisions during embryonic development [38]. Both the magnitude and duration of WNT signaling influence primitive streak progression, with increasing cumulative WNT exposure progressively favoring definitive endoderm (DE), lateral plate mesoderm (LPM), and paraxial mesoderm (PM) fates [21,37], with CMs arising from a narrow LPM-favoring window. This, in turn, constrains the range of signaling conditions permissive for efficient CM differentiation, such that even small variations in WNT activity may divert cells toward alternative lineages, leading to variable outcomes. Thus, a process design that reshapes WNT-dependent lineage boundaries to broaden the cardiac-permissive window may be key to achieving robust CM differentiation.

The GiWi protocol was originally developed with 1-day CHIR treatment followed by WNT inhibition from day 3 of differentiation [14,15], with some alternative protocols employing 2-day CHIR treatment and WNT inhibition initiated on day 2 [12,16,39]. Meanwhile, in a systematic study of developmental lineage decisions during PSC differentiation, Loh *et al*. [40] reported that, following 1-day treatment with a CHIR-containing cocktail for primitive streak induction, WNT inhibition initiated as early as day 1 suppresses PM specification while promoting LPM differentiation. However, the effect of earlier WNT inhibition on CM differentiation has remained controversial, with previous studies reporting either enhanced [17,25] or reduced [15,21] differentiation. Nevertheless, given the WNT-dependent patterning from DE through LPM to PM fates [21,37], we reasoned that early WNT inhibition could promote LPM specification under conditions of strong initial WNT activation resulting from high CHIR concentration and/or high cellular responsiveness to CHIR, which would otherwise favor PM specification. We further hypothesized that this altered lineage preference could extend the effective initial WNT activation range for CM differentiation, thereby improving its robustness through process window expansion.

In the present study, we systematically examined the effects of early WNT inhibition initiated on day 1 on the effective CHIR concentration window and further expanded our analysis to a two-dimensional process window by incorporating cell density as an additional parameter. Additionally, we analyzed the temporal expression of lineage-specific transcription factors (TFs), including TFs associated with cardiac and competing lineages, to characterize the expression patterns underlying process window expansion and improved differentiation efficiency. Finally, we evaluated process robustness by comparing the early WNT inhibition protocol with the standard protocol under parameter perturbations and across wells and batches, followed by assessment of the phenotypic properties of the resulting CMs. This study provides, to our knowledge, the first demonstration that early WNT inhibition, guided by developmental principles of lineage specification, can substantially expand the multidimensional process window of CM differentiation and thereby improve process robustness.

## 2. Materials and methods

### 2.1. hiPSC maintenance

The hiPSC lines 610B1 (Cell line ID: HPS0331) [41] and 253G1 (Cell line ID: HPS0002) [42] were provided by the RIKEN BioResource Research Center (RIKEN BRC) through the National BioResource Project of the Ministry of Education, Culture, Sports, Science and Technology, Japan. The hiPSCs were maintained in mTeSR Plus medium (STEMCELL Technologies, Vancouver, BC, Canada; Cat. # 100-0276) on culture dishes coated with hESC-qualified Matrigel (Corning, New York, NY, USA; Cat. # 354277). Matrigel was diluted in cold DMEM/F12 (Nacalai Tesque, Kyoto, Japan; Cat. # 11583-95) according to the manufacturer’s recommended dilution, and the dishes were coated overnight at 4 °C. The culture medium was changed every 2 days. The cells were passaged as clumps every 3–4 days using 0.5 mM ethylenediaminetetraacetic acid (EDTA) with split ratios of 1:8–1:16.

### 2.2. Cardiac differentiation

Cardiac differentiation was performed by biphasic WNT signaling modulation. CHIR (Cayman Chemical, Ann Arbor, MI, USA; Cat. # 13122) was administered from days 0 to 1 of differentiation, followed by IWP-2 treatment (Selleck Chemicals, Houston, TX, USA; Cat. # S7085) from days 3 to 5 for the control condition, as described by Lian *et al*. [14,15], or from days 1 to 5 for the test condition.

Three days before the initiation of differentiation, hiPSCs were dissociated with Accutase (Nacalai Tesque; Cat. # 12679-54) and seeded into a 48-well plate (Greiner Bio-One, Kremsmunster, Austria; Cat. # 677180) coated with Matrigel in mTeSR Plus medium supplemented with 10 μM Y-27632 (Kanto Chemical, Tokyo, Japan; Cat. # A3008-10) for the first 24 h. The cell-seeding density was varied depending on the experiment, as specified below. Cells were expanded for 3 days with daily replacement of mTeSR Plus medium. On day 0 of differentiation, the medium was replaced with differentiation medium consisting of RPMI 1640 (RPMI; Nacalai Tesque; Cat. # 30264-85) supplemented with 2% B27 minus insulin (Thermo Fisher Scientific, Waltham, MA, USA; Cat. # A1895601), hereafter referred to as RPMI/B27 (−ins), and CHIR at the concentration specified for each experiment. On day 1, exactly 24 h after differentiation was initiated, the medium was replaced with RPMI/B27 (−ins) without IWP-2 for the control condition or with 5 μM IWP-2 for the test condition. On day 3, the medium was replaced with RPMI/B27 (−ins) containing 5 μM IWP-2, followed by replacement with fresh RPMI/B27 (−ins) on day 5 to remove IWP-2. From day 7 onward, the medium was replaced with RPMI supplemented with 2% B27 containing insulin (Thermo Fisher Scientific; Cat. # 17504044), hereafter referred to as RPMI/B27 (+ins). On day 10, cells were harvested using TrypLE Express (Thermo Fisher Scientific; Cat. # 12604021) to measure the percentage of cardiac troponin T (cTNT)-positive cells by flow cytometry or for downstream analyses, as described below.

In the initial experiment assessing whether the early WNT inhibition expanded the effective CHIR concentration window, CHIR was varied from 4.5 to 13.5 μM for the 610B1 cell line and from 6 to 20 μM for the 253G1 cell line, while the cell-seeding densities were fixed at 2.2 × 10^4^ and 1.6 × 10^4^ cells/cm^2^, respectively. For the two-dimensional evaluation of the process window defined by CHIR concentration and cell-seeding density using the 610B1 cell line, the two parameters were varied from 5.5 to 13.5 μM and from 1.1 to 4.4 × 10^4^ cells/cm^2^, respectively.

To test the effect of WNT inhibition duration using the 610B1 cell line, the medium was replaced with RPMI/B27 (−ins) supplemented with 5 μM IWP-2 on day 1, and IWP-2 treatment was continued for 2, 3, or 4 days. To test the effect of different WNT inhibitors using the 610B1 cell line, the medium was replaced with RPMI/B27 (−ins) supplemented with 5 μM IWP-2, 2 μM Wnt-C59 (C59; Cayman Chemical; Cat. # 16644), or 10 μM XAV-939 (XAV; Selleck Chemicals; Cat. # S1180) on day 1, and the treatment was continued for 4 days. For these experiments, CHIR concentration and cell-seeding density were set at 10.5 μM and 2.2 × 10^4^ cells/cm^2^, respectively. A control differentiation in which IWP-2 was administered from days 3 to 5 was conducted in parallel. After WNT inhibitor withdrawal, the cells were cultured according to the procedure described above.

Process robustness was assessed using the 610B1 cell line. To assess robustness to parameter perturbations, nine conditions were tested, comprising the optimal condition (shown in the corresponding figure) and conditions with CHIR concentrations shifted by ±10% and cell-seeding densities shifted by ±15% relative to the corresponding optimal values for each protocol. Well-to-well and batch-to-batch variability in differentiation efficiency was assessed by repeatedly evaluating the optimal condition across wells and independent experimental batches.

### 2.3. Long-term CM culture

On day 10 of differentiation, cells were dissociated using TrypLE Express and replated into a 12-well plate (AGC Techno Glass, Shizuoka, Japan; Cat. # 3815-012) coated with type I collagen (Nitta Gelatin, Osaka, Japan; Cat. # 631-00771) to double the culture area, using RPMI supplemented with 10% fetal bovine serum (FBS; Sigma-Aldrich, St. Louis, MO, USA; Cat. # 172012), 1% penicillin–streptomycin (PS; Fujifilm Wako Pure Chemical, Osaka, Japan; Cat. # 168-23191), and 10 μM Y-27632. From days 11 to 15, the cells were subjected to metabolic selection [29,30] in glucose-free RPMI (Nacalai Tesque; Cat. # 09892-15) supplemented with 4 mM L-lactate (Fujifilm Wako Pure Chemical; Cat. # 129-02666), 213 μg/mL L-ascorbic acid 2-phosphate (Sigma-Aldrich; Cat. # A8960), 0.25% bovine albumin fraction V (Thermo Fisher Scientific; Cat. # 15260037), and 1% PS, with the medium refreshed on day 13. On day 15, cells were dissociated using TrypLE Express and replated into a 48-well plate (AGC Techno Glass; Cat. # 3860-048) or 96-well plate (AGC Techno Glass; Cat. # 3860-096), depending on the downstream analysis, at 7.6 × 10^4^ cells/cm^2^, a density selected to achieve confluence, in RPMI supplemented with 10% FBS, 1% PS, and 10 μM Y-27632. From day 16 onward, the cells were cultured in RPMI/B27 (+ins) supplemented with 1% PS, with the medium replaced every other day. On day 30, the cells were subjected to flow cytometry, gene expression analysis, and calcium transient analysis, as described below.

### 2.4. Generalized additive model (GAM) analysis

We used GAMs [43,44] to characterize nonlinear responses. Cardiac differentiation efficiency or the absolute difference in differentiation efficiency between batches was expressed as a proportion and logit-transformed prior to model fitting, as shown below:

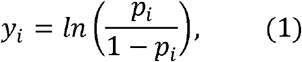

where *p_i_* denotes the differentiation efficiency or the absolute difference between batches, expressed as a proportion, for observation *i*, and *y_i_* denotes its logit-transformed value.

For analyses using CHIR concentration as the only explanatory variable, a GAM was specified as follows:

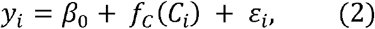

where *β_i_* is the intercept, *C_i_* is the CHIR concentration, *f_c_*(·) is a smooth function represented by cubic P-splines, and *ε_i_* is the residual error.

For analyses using both CHIR concentration and cell-seeding density as explanatory variables, a GAM was specified as follows:

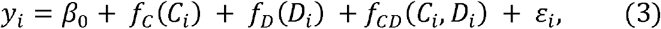

where *D_i_* is the cell-seeding density, *f_C_*(·) and *f_D_*(·) are smooth functions represented by cubic P-splines, and *f_CD_*(·, ·) is a tensor-product smooth representing the interaction between CHIR concentration and cell-seeding density.

The models were fitted using LinearGAM implemented in the pyGAM package (v0.10.1) [45] in Python (v3.9.23). The smoothing parameter was selected by grid search over logarithmically spaced candidate values ranging from 10^−3^ to 10^3^ by minimizing the generalized cross-validation score. The number of spline basis functions was selected for each analysis based on model adequacy as well as visual agreement between observed data and fitted curves. Model adequacy was assessed using predicted-versus-observed, residual-versus-predicted, and normal quantile–quantile plots of the residuals. Predicted values were back-transformed to the original scale using the inverse-logit transformation for visualization.

### 2.5. Measurement of cell confluency

Cell confluency was measured by image analysis using Trainable Weka Segmentation [46], a plugin for Fiji (ImageJ2 v2.16.0; National Institutes of Health, Bethesda, MD, USA) [47], which performs pixel-based classification using machine learning. On day 0 of differentiation, phase-contrast images from five representative wells were obtained for each condition. A representative image was used to annotate cell and non-cell areas and train a classification model using Trainable Weka Segmentation. The model was applied to all images to generate binary images distinguishing cell and non-cell areas. Small noise objects in non-cell regions were removed using the particle detection function, and small holes within cell areas were filled. Minor erosion and dilation adjustments were applied to better match the binary images to the cell-covered areas. Confluency was then calculated by dividing the number of pixels classified as cell areas by the total number of pixels in the image and multiplying the resulting value by 100. Post-processing steps following Trainable Weka Segmentation were scripted in Fiji to ensure consistent processing across all images.

### 2.6. Flow cytometry

Cells were stained with the LIVE/DEAD Fixable Violet Dead Cell Stain Kit (Thermo Fisher Scientific; Cat. # L34963) following the manufacturer’s protocol prior to fixation and fluorescence staining. On day 10, cells were fixed and permeabilized using the Foxp3/Transcription Factor Staining Buffer Set (Thermo Fisher Scientific; Cat. # 00-5523-00) in accordance with the manufacturer’s instructions. Cells were then stained at 4 °C for 30 min with Alexa Fluor 647–conjugated anti-cTNT antibody (1:50 dilution; BD Biosciences, San Jose, CA, USA; Cat. # 565744), followed by washing with phosphate-buffered saline (PBS) containing 2% FBS and 0.5 mM EDTA, hereafter referred to as staining buffer. On day 30 of differentiation, cells were fixed with 2% paraformaldehyde solution at room temperature (20–25 °C) for 10 min, followed by washing with PBS. Cells were then permeabilized with prechilled 100% methanol on ice for 15 min, followed by washing with staining buffer. Next, the cells were co-stained at room temperature for 30 min with the same Alexa Fluor 647–conjugated anti-cTNT antibody (1:50 dilution), Vio B515–conjugated recombinant human anti-MLC2a antibody (1:50 dilution; Miltenyi Biotec, Bergisch Gladbach, Germany; Cat. # 130-129-272), and PE-conjugated recombinant human anti-MLC2v antibody (1:50 dilution; Miltenyi Biotec; Cat. # 130-119-680), followed by washing with staining buffer. The stained samples were analyzed using a BD FACSCanto II flow cytometer (BD Biosciences). The data were analyzed using FlowJo software (version 10.10; BD Biosciences). For day 10 cells, gating thresholds were determined based on the distribution of the cTNT-negative population in a low-differentiation sample. For day 30 cells, thresholds were determined based on the distributions of fluorescence-minus-one (FMO) controls.

### 2.7. Quantitative polymerase chain reaction (qPCR) analysis

RNA was extracted using the RNeasy Micro Kit (QIAGEN, Venlo, Netherlands; Cat. # 74004) following the manufacturer’s instructions. For each sample, 100−500 ng of total RNA was reverse-transcribed into cDNA using the ReverTra Ace qPCR RT Master Mix with gDNA Remover (Toyobo, Osaka, Japan; Cat. # FSQ-301) at 37 °C for 15 min, followed by 50 °C for 5 min. qPCR was conducted using cDNA corresponding to 3 ng of input RNA as the template with the SYBR Green qPCR Master Mix (Thermo Fisher Scientific; Cat. # A66732) on the StepOne Real-Time PCR System (Thermo Fisher Scientific). The thermal cycling conditions consisted of an initial denaturation at 95 °C for 2 min, followed by 40 cycles of denaturation at 95 °C for 15 s and annealing and extension at 60 °C for 1 min. The primers used in this study are listed in Table S1.

### 2.8. RNA sequencing (RNA-seq) and data processing

RNA was extracted as described above. mRNA was isolated using the NEBNext Poly(A) mRNA Magnetic Isolation Module (New England Biolabs, Ipswich, MA, USA; Cat. # E7490), and sequencing libraries were prepared using the NEBNext Ultra II RNA Library Prep Kit for Illumina (New England Biolabs; Cat. # E7770) according to the manufacturer’s instructions. Sequencing was performed on a NextSeq 550 system (Illumina, San Diego, CA, USA) using the NextSeq 500/550 High Output Kit v2.5 (Illumina; Cat. # 20024906) with 81-bp single-end reads. Raw sequencing reads were trimmed using BBDuk implemented in BBTools (v39.88) [48]. Read quality was assessed using FastQC (v0.12.1; Babraham Bioinformatics) before and after trimming. The processed reads were aligned to the human reference genome GRCh38.p14 with the GENCODE release 49 gene annotation [49] using STAR (v2.7.11b) [50]. Gene count data were generated using RSEM (v1.3.1) [51]. Count normalization and differential expression analysis were performed using DESeq2 (v1.50.2) [52] in R (v4.5.2; R Core Team, 2025). A list of cardiac-enriched genes (CEGs) was obtained from the Human Protein Atlas database [53,54], as described in the Supplementary Methods.

### 2.9. Calcium transient analysis

On day 30, confluent cells prepared as described above in the 96-well plate were loaded with Fluo-4 using the Calcium Kit II (Dojindo, Kumamoto, Japan; Cat. # CS32) according to the manufacturer’s instructions. Two platinum wire electrodes were positioned 5 mm apart, parallel to and in close proximity to the cell layer. Electrical pulse stimulation (EPS; 4 V/mm, 0.5 Hz, 10 ms pulse duration) was applied using an EPS generator (C-PACE EP; IonOptix, Westwood, MA, USA) under a fluorescence microscope (IX81; Olympus, Tokyo, Japan) equipped with a high-speed camera (HAS-U1M; DITECT, Tokyo, Japan). Videos were recorded from 6 to 9 fields of view for each sample at 100 frames per second with an exposure time of 10 ms. For each video, the mean fluorescence intensity across the entire field of view was extracted over time using OpenCV (v4.11.0) [55] in Python, and the ΔF/F_0_ waveform was calculated. For each field of view, the peak value was calculated as the mean peak value of three consecutive calcium transients. The peak ΔF/F_0_ values obtained from the 6 to 9 fields of view were then averaged, and this mean value was used as the representative peak ΔF/F_0_ value for each sample.

### 2.10. Statistical analysis

For comparisons between two groups, parametric or nonparametric tests were selected as appropriate. Welch’s *t*-test and the paired *t*-test were used for parametric unpaired and paired comparisons, respectively, whereas the Mann–Whitney U and Wilcoxon signed-rank tests were used for nonparametric unpaired and paired comparisons, respectively. Differences in dispersion were assessed using Levene’s test. Multiple-comparison correction was applied as appropriate using the Benjamini–Hochberg (BH) method. For comparisons among multiple groups, one-way analysis of variance (ANOVA) was performed, followed by Tukey’s post hoc test. Data are presented as the mean ± standard deviation (SD), median with the interquartile range (IQR), or box plots with optional violin overlays, as appropriate. The sample size (*n*) and type of replicate are indicated in each figure legend. A *p*-value < 0.05, or an adjusted *p*-value < 0.05 when applicable, was considered statistically significant. For RNA-seq data analysis, genes showing an adjusted *p*-value < 0.05 and absolute log_2_ fold change ≥ 1 were considered differentially expressed. Statistical analyses and data visualization were performed using R or Python.

## 3. Results

### 3.1. Early WNT inhibition expands the CHIR concentration window that allows efficient CM differentiation

Based on our hypothesis that early WNT inhibition could extend the effective initial WNT activation range by suppressing PM specification, we first examined whether it broadened the CHIR concentration window that allows high-efficiency CM differentiation. Therefore, we compared the widely used standard GiWi (S-GiWi) protocol developed by Lian *et al*. [14,15], in which WNT inhibition was applied from days 3 to 5, with a modified GiWi protocol in which WNT inhibition was initiated on day 1 and continued through day 5 (Fig. 1A). We employed two iPSC lines, 610B1 and 253G1, which differ in their tissue of origin and reprogramming method, to examine the cell line dependence of this effect. We modeled the nonlinear relationship between CHIR concentration and CM differentiation efficiency using GAMs [43,44], which provide greater flexibility for capturing complex nonlinear responses across a broad parameter range than conventional low-order polynomial models commonly used in design of experiment–based process development [22,56,57]. In the 610B1 cell line, the modified GiWi protocol yielded slightly lower differentiation efficiency than that achieved with S-GiWi at 7.5 μM CHIR (77.7 ± 4.5% for S-GiWi vs. 66.6 ± 4.9% for modified GiWi), whereas this relationship was reversed at 9.0 μM, with modified GiWi yielding slightly higher efficiency than that achieved with S-GiWi (85.0 ± 6.5% for S-GiWi vs. 92.9 ± 1.6% for modified GiWi) (Fig. 1B and C). At concentrations of 10.5 μM or higher within the tested range, the modified GiWi maintained significantly higher efficiencies, whereas S-GiWi resulted in little cardiac differentiation. Consistent with the concentration–response analysis, the modified GiWi protocol increased the width of the ≥80% window by 2.0-fold (Fig. 1D) and ≥80% AUC normalized to the window-center concentration by 2.8-fold (Fig. 1E) compared with those obtained with S-GiWi. Similarly, in the 253G1 cell line, the modified GiWi yielded lower differentiation efficiencies at concentrations of 12 μM or lower but maintained significantly higher efficiencies at 14 μM or higher (Fig. 1F and G). It increased the width of the ≥80% window by 1.7-fold (Fig. 1H) and normalized ≥80% AUC by 3.3-fold (Fig. 1I). Thus, the expansion of the effective CHIR concentration window toward higher concentrations was reproduced in both cell lines, demonstrating that early WNT inhibition broadens this window independent of iPSC line.

**Fig. 1.**
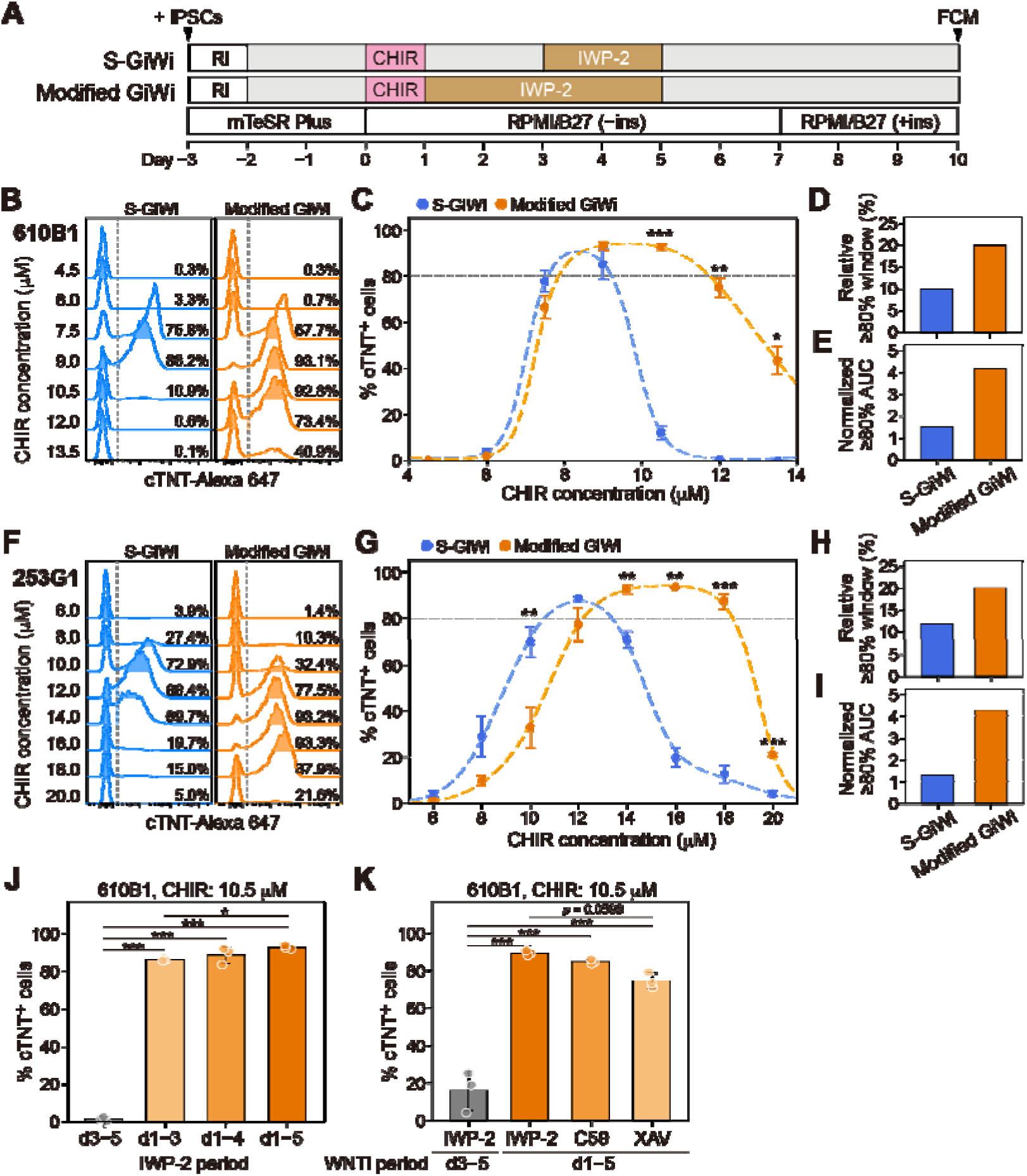
Effect of early WNT inhibition on expansion of the CHIR concentration window enabling high-efficiency cardiac differentiation. **A:** Schematic of the S-GiWi and modified GiWi protocols, with IWP-2 treatment applied on days 3–5 and 1–5, respectively. Ri, ROCK inhibitor (Y-27632); FCM, flow cytometry. **B:** Representative flow cytometry histograms for cTNT expression on day 10 of differentiation in the 610B1 cell line with 4.5−13.5 μM CHIR. **C:** Percentages of cTNT-positive cells on day 10 in the 610B1 cell line with 4.5−13.5 μM CHIR. The data were fitted using GAMs; the fitted curves are shown as dashed lines. Statistical analyses were performed using Welch’s *t*-test to compare the data at each CHIR concentration, followed by BH correction. **D:** Relative CHIR concentration window yielding ≥80% cTNT-positive cells around the center concentration of the window, calculated from the fitted data shown in (C). **E:** Area under each fitted curve shown in (C) above the 80% threshold (≥80% AUC), normalized by dividing by the corresponding window center concentration. **F−I:** Corresponding analyses using the 253G1 cell line with 6.0−20.0 μM CHIR. **J:** Percentages of cTNT-positive cells on day 10 in the 610B1 cell line using 10.5 μM CHIR with IWP-2 treatment applied for different durations: days 3−5, 1−3, 1−4, or 1−5. **K:** Percentages of cTNT-positive cells on day 10 in the 610B1 cell line using 10.5 μM CHIR with IWP-2 treatment from days 3 to 5 or with IWP-2, C59, or XAV treatment from days 1 to 5. WNTi, WNT inhibitor. Statistical analyses in (J) and (K) were performed using one-way ANOVA followed by Tukey’s post hoc test. Data in (C), (G), (J), and (K) are presented as mean ± SD (n = 3 wells). \**p* < 0.05, \*\**p* < 0.01, and \*\*\**p* < 0.001.

In addition to the GiWi protocol established by Lian *et al*., another treatment schedule involving 2 days of CHIR treatment followed immediately by 2 days of WNT inhibition has also been widely used with B27-based media [16,39], including in this study, as well as with CDM3 medium [58]. To determine the effective CHIR concentration window under this schedule, we examined the CHIR dose–response relationship using the 610B1 cell line (Fig. S1). Cardiac differentiation efficiency exceeded 80% only within a narrow range of 6.0–6.4 μM and sharply decreased above 6.5 μM, resulting in a narrower window than that observed with the modified GiWi protocol. These results suggest that expansion of the effective CHIR concentration window may depend on the use of a 1-day CHIR treatment followed immediately by early WNT inhibition.

Next, we conducted two independent experiments to examine whether the duration and pharmacological mechanism of early WNT inhibition influenced this effect, using 10.5 μM CHIR, at which S-GiWi yielded a low CM proportion in the 610B1 cell line (Fig. 1C). Initiating IWP-2 treatment on day 1 markedly increased differentiation efficiency compared with that achieved with the standard day 3–5 treatment, even when treatment was limited to days 1–3 (Fig. 1J). Extending IWP-2 treatment to days 1–4 and 1–5 further increased differentiation efficiency in a duration-dependent manner. Subsequently, we compared IWP-2 with C59, another porcupine inhibitor that blocks WNT ligand secretion, and XAV, a tankyrase inhibitor that stabilizes AXIN and promotes β-catenin degradation. When administered from days 1–5, all three inhibitors markedly increased differentiation efficiency relative to that achieved with the standard day 3–5 IWP-2 treatment (Fig. 1K). C59 produced an effect closer to that of IWP-2, whereas XAV produced a somewhat weaker effect. Collectively, these results indicate that a longer duration of early WNT inhibition is more effective and that the porcupine inhibitors tested are more effective than the tankyrase inhibitor tested in this context.

Given its ability to broaden the effective CHIR concentration window, we hereafter refer to the modified GiWi protocol with IWP-2 treatment from days 1 to 5 as the Robust GiWi (R-GiWi) protocol.

### 3.2. Early WNT inhibition–based process suppresses key TFs associated with DE and PM programs

Given the substantial expansion of the effective CHIR concentration window toward higher concentrations, the optimal operating point for R-GiWi shifted toward a higher concentration than that for S-GiWi. This led us to hypothesize that R-GiWi at a higher setpoint would more effectively suppress DE specification, which is favored by lower levels of WNT signaling. Furthermore, although the higher-range expansion of the effective CHIR concentration window was likely to involve suppression of PM specification, the expression patterns of PM-associated TFs remained uncharacterized. To address these two issues and more broadly capture lineage-associated transcriptional features, we analyzed the temporal expression of a panel of TFs associated with early developmental programs under conditions selected from the CHIR concentration–response curves in Fig. 1C. Specifically, the conditions were S-GiWi with 8.5 μM CHIR, representing the approximate optimal concentration identified at this stage; S-GiWi with 10.5 μM CHIR, a high concentration at which cardiac differentiation was markedly reduced; and R-GiWi with 10.5 μM CHIR, which maintained efficient cardiac differentiation and was near the center of the ≥80% window (Fig. 2A). These conditions are hereafter referred to as S-GiWi (C8.5), S-GiWi (C10.5), and R-GiWi (C10.5), respectively.

**Fig. 2.**
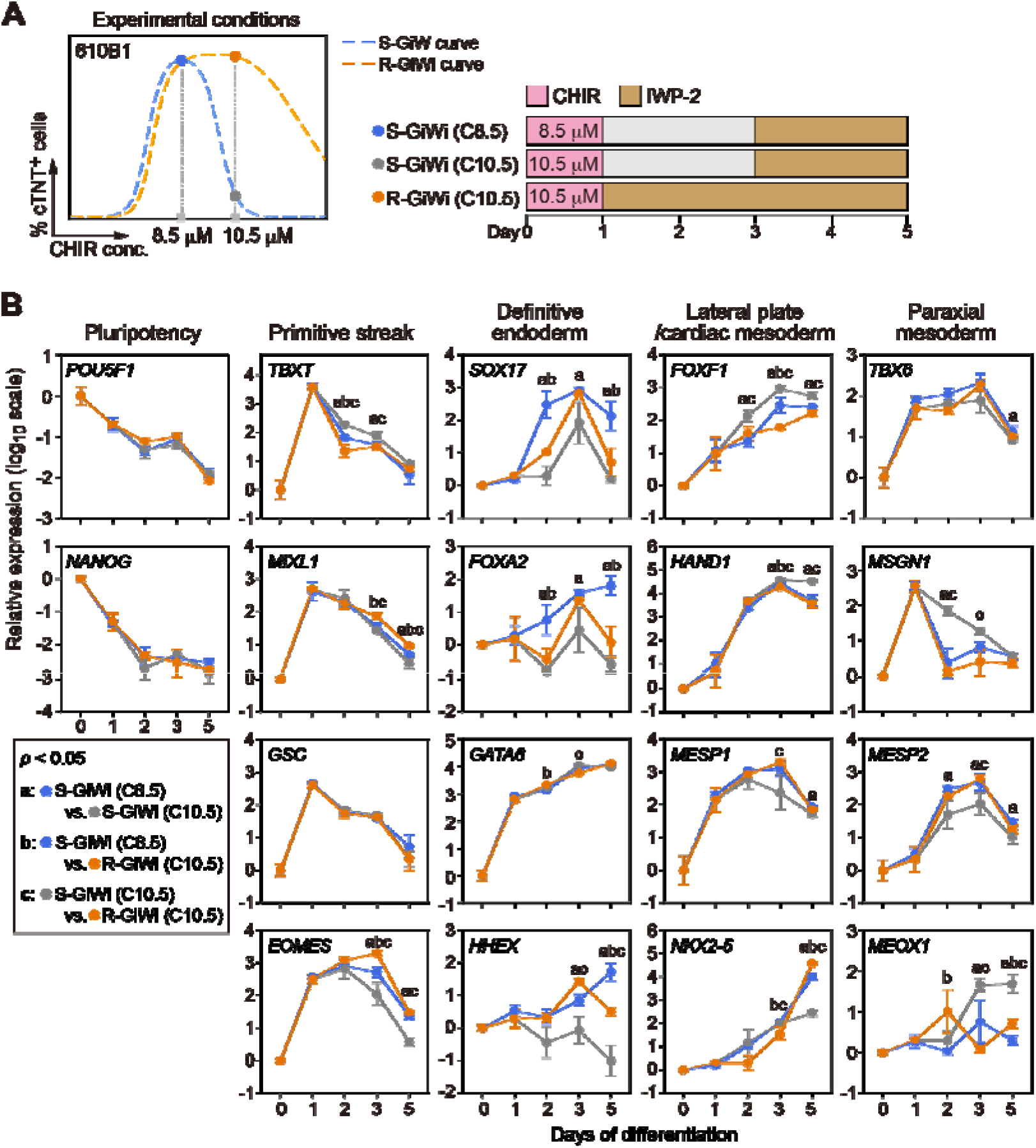
Effects of early WNT inhibition on lineage-associated TF expression. **A:** Experimental conditions used for TF expression analysis. The differentiation efficiency curves shown on the left correspond to those obtained with the 610B1 cell line in Fig. 1C. Three conditions were selected: S-GiWi with 8.5 μM CHIR (blue circle), S-GiWi with 10.5 μM CHIR (gray circle), and R-GiWi with 10.5 μM CHIR (orange circle), referred to as S-GiWi (C8.5), S-GiWi (C10.5), and R-GiWi (C10.5), respectively. **B:** Expression profiles of lineage-associated TFs on days 0, 1, 2, 3, and 5 of differentiation. Data are presented as mean ± SD (n = 3 wells). Statistical analyses were performed on days 2, 3, and 5 using one-way ANOVA followed by Tukey’s post hoc test. Letters indicate statistically significant differences (*p* < 0.05): a, S-GiWi (C8.5) versus S-GiWi (C10.5); b, S-GiWi (C8.5) versus R-GiWi (C10.5); and c, S-GiWi (C10.5) versus R-GiWi (C10.5). Expression profiles of additional lineage-associated markers are provided in Fig. S2.

Regarding the hypothesis concerning DE suppression, we focused, for simplicity, on the comparison between S-GiWi (C8.5) and R-GiWi (C10.5). As shown in the DE TF panel of Fig. 2B, R-GiWi (C10.5) significantly suppressed the expression of *SOX17* and *FOXA2*, two major early TFs involved in the DE program [59], on days 2 and 5 compared with that under S-GiWi (C8.5). Although *GATA6* expression was largely maintained, this did not necessarily indicate preserved DE specification because *GATA6* also contributes to precardiac mesoderm development [60,61]. R-GiWi (C10.5) also significantly reduced the expression of *HHEX*, a marker for anterior endoderm [62], on day 5. These results strongly suggest that R-GiWi suppressed both initial DE specification and subsequent progression toward the anterior endoderm lineage. In this experiment, cardiac differentiation efficiencies were 85.3% for S-GiWi (C8.5) and 90.2% for R-GiWi (C10.5), both close to the earlier results (Fig. 1C), with the modestly higher efficiency in R-GiWi possibly reflecting DE suppression.

Next, we focused on the comparison between S-GiWi (C10.5) and R-GiWi (C10.5) to characterize the expression patterns of PM-associated TFs underlying the higher-range expansion of the CHIR concentration window. In this experiment, cardiac differentiation efficiency was only 11.0% for S-GiWi (C10.5), which closely matches the earlier result (Fig. 1C) but is markedly lower than the 90.2% achieved with R-GiWi (C10.5). As shown in the PM TF panel of Fig. 2B, R-GiWi (C10.5) had little effect on *TBX6* expression but sharply suppressed *MSGN1* expression immediately after the initiation of WNT inhibition, resulting in approximately 60-fold lower expression than that in S-GiWi (C10.5) on day 2. *TBX6* and *MSGN1* are key regulators of the initial PM-specification program [63–67]; thus, early WNT inhibition substantially impaired a key component of this program. *MESP2*, a regulator associated with presomitic mesoderm segmentation, was expressed at higher levels in R-GiWi (C10.5) despite its much higher cardiac differentiation efficiency. The higher *MESP2* expression was consistent with the preserved and slightly elevated expression of *TBX6*, an upstream transcriptional regulator of *MESP2* [68]. The higher expression of *DLL1* in R-GiWi (Fig. S2), another downstream target of *TBX6* [69], further supported preservation of this axis. In contrast, *CDX2*, a regulator associated with posterior PM identity, was reduced in R-GiWi (C10.5) on days 2 and 3 (Fig. S2). *MEOX1*, a regulator of subsequent somitic mesoderm development, was also significantly reduced on days 3 and 5, suggesting that early WNT inhibition suppressed the posterior PM program and its subsequent progression. Collectively, although early WNT inhibition did not uniformly suppress PM-associated genes, the marked suppression of *MSGN1* emerged as a key transcriptional feature associated with the higher-range expansion of the CHIR concentration window, potentially contributing to this effect through altered lineage preference away from PM and toward cardiac fates.

Notably, *FOXF1*, a regulator of LPM development, was expressed at lower levels in R-GiWi (C10.5) than in both S-GiWi (C8.5) and S-GiWi (C10.5), with the difference being pronounced on day 3 (Fig. 2B), showing an expression pattern opposite to the trend of CM differentiation efficiency. This reduction was accompanied by a tendency toward delayed induction of *NKX2-5*, a regulator of cardiac mesoderm development, on days 2 and 3 in R-GiWi (C10.5). *HAND1* and *MESP1* expression trends were comparable between S-GiWi (C8.5) and R-GiWi (C10.5). These findings suggest that early WNT inhibition did not directly promote the LPM/cardiac mesoderm program. Rather, the expansion of the CHIR concentration window along with improved differentiation efficiency with R-GiWi (C10.5) reflects an altered lineage preference resulting from the suppression of competing DE and PM programs. Moreover, the overall transcriptional features suggest that R-GiWi may be less susceptible to deviation toward competing fates.

### 3.3. Early WNT inhibition substantially expands the multidimensional process window

Next, we characterized the two-dimensional process windows defined by CHIR concentration and cell density, another critical parameter affecting differentiation efficiency [37,70–72], for the S-GiWi and R-GiWi protocols. For this purpose, we conducted two independent experiments, with both protocols performed in parallel within each experiment across all 25 combinations of five levels of each factor to model the differentiation responses.

Cell confluency at the onset of differentiation, rather than the initial cell-seeding density itself, is considered more relevant to differentiation outcomes, as it more directly reflects the cellular states immediately before differentiation [19] and paracrine environment during differentiation [37], both of which can influence cellular responsiveness to CHIR. Thus, we quantified confluency at the onset of differentiation for each cell-seeding density using machine learning–based image analysis (Fig. 3A). Although statistically significant differences in confluency were observed between the two experiments at two seeding densities, the differences were modest, and the overall relationship between cell-seeding density and confluency remained similar across experiments (Fig. 3B).

**Fig. 3.**
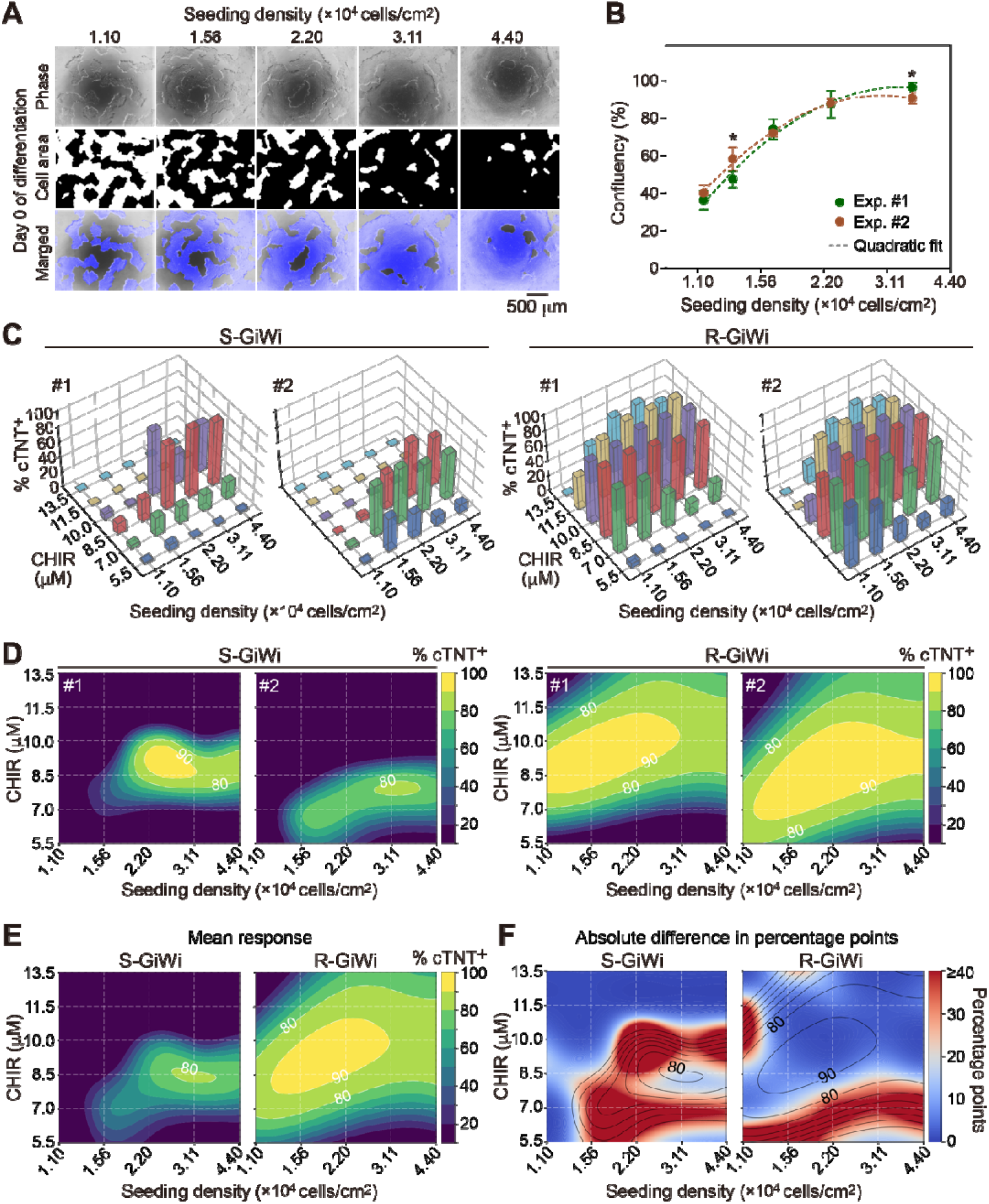
Effects of early WNT inhibition on the two-dimensional process window defined by CHIR concentration and cell-seeding density. **A:** Representative phase-contrast images at the onset of differentiation (top), segmented cell-area images in which cell areas are highlighted in black (middle), and merged images with segmented cell areas shown in blue (bottom) for 610B1 cells seeded at the indicated densities 3 days before differentiation. Scale bar, 500 μm. **B:** Cell confluency on day 0 at each cell-seeding density in two independent experiments. The dashed curves represent quadratic fits to the data from each experiment. Data are presented as mean ± SD (n = 5 wells per cell-seeding condition). Statistical analyses were performed using Welch’s *t*-test to compare the two experiments at each cell-seeding density, followed by BH correction. \**p* < 0.05. **C:** Percentages of cTNT-positive cells on day 10 obtained using S-GiWi or R-GiWi across combinations of CHIR concentration (5.5−13.5 μM) and cell-seeding density (1.1−4.4 × 10^4^ cells/cm^2^) in two independent experiments (n = 1 well per condition in each experiment). **D:** Contour plots of the GAM-fitted response surfaces for each experiment. **E:** Contour plots of the GAM-fitted response surfaces based on the mean responses of the two experiments. **F:** Contour plots of absolute differences between the two experiments modeled using GAMs, expressed in percentage points. Black contour lines represent the GAM-fitted mean response surfaces shown in (E). Model adequacy assessments corresponding to this figure are presented in Figs. S3 and S4.

The differentiation efficiencies shown in Fig. 3C were used to fit separate GAMs for each protocol and experiment, yielding the response surfaces shown in Fig. 3D. R-GiWi consistently yielded a substantially broader ≥80% window than that of S-GiWi in both experiments, with a large ≥90% region also reproducibly observed. In contrast, S-GiWi not only yielded narrower windows but also exhibited noticeable shifts in the ≥80% window and response peak between the two experiments, despite similar overall confluency on day 0, as observed earlier. This suggests that consistently achieving high efficiency at a fixed operating point may be highly challenging with S-GiWi. The high-efficiency windows of R-GiWi also shifted between experiments; however, their greater breadth may reduce the practical impact of these shifts on differentiation consistency at a fixed operating point.

Next, we modeled the differentiation responses using the mean efficiencies from the two experiments to derive high-efficiency process windows representative of their combined responses (Fig. 3E). This enables a more reliable prediction of differentiation efficiencies across the parameter space and facilitating the subsequent selection of operating conditions. Averaging the responses from the two experiments clearly highlighted the substantially larger ≥80% window for R-GiWi relative to that for S-GiWi than the individual-experiment analyses did (Figs. 1C and 3D). Furthermore, mapping the modeled absolute differences in percentage points between the two experiments onto the averaged response surfaces revealed that the central region of the high-efficiency window for R-GiWi was farther from regions showing large between-experiment differences than the corresponding region for S-GiWi (Fig. 3F). Collectively, these results suggest that operating within the central region of this window with R-GiWi would provide greater robustness to variations in CHIR concentration and cell density, as well as differences in cellular responsiveness across wells and batches.

### 3.4. Early WNT inhibition–based process shows substantially greater robustness

To validate the robustness of R-GiWi, we first investigated the effects of perturbations in CHIR concentration and cell density on differentiation efficiency. This evaluation was designed to assess sensitivity to controlled changes in these factors but also to provide a predictive surrogate assessment of process robustness against well-to-well and batch-to-batch variability. Such variation may arise from differences in cellular responsiveness to WNT activation cues and paracrine signaling environments, as well as from technical variation affecting these critical process parameters. For this purpose, we set nine experimental conditions: the approximate center of the high-efficiency window (the ≥80% window for S-GiWi and ≥90% window for R-GiWi) as the predicted optimal condition (black circle), and eight additional conditions (colored circles) representing feasible deviations that could reflect both biological and technical variations (Fig. 4A). The latter conditions were generated by perturbing the CHIR concentration by ±10% and cell-seeding density by ±15% relative to the corresponding values in the predicted optimal condition. Then, we compared the observed efficiencies of these nine conditions between S-GiWi and R-GiWi, with the corresponding model predictions also shown for reference (Fig. 4B). Although the observed efficiencies were more variable than the model predictions, reflecting the limited ability of the models to capture the inherent variability, the relative differences in both differentiation efficiency and variability between R-GiWi and S-GiWi were consistent with the model predictions. R-GiWi achieved significantly higher differentiation efficiencies and lower variability than those achieved with S-GiWi, with efficiencies exceeding 85% across all R-GiWi conditions. These results demonstrate markedly greater robustness of R-GiWi to parameter perturbations.

**Fig. 4.**
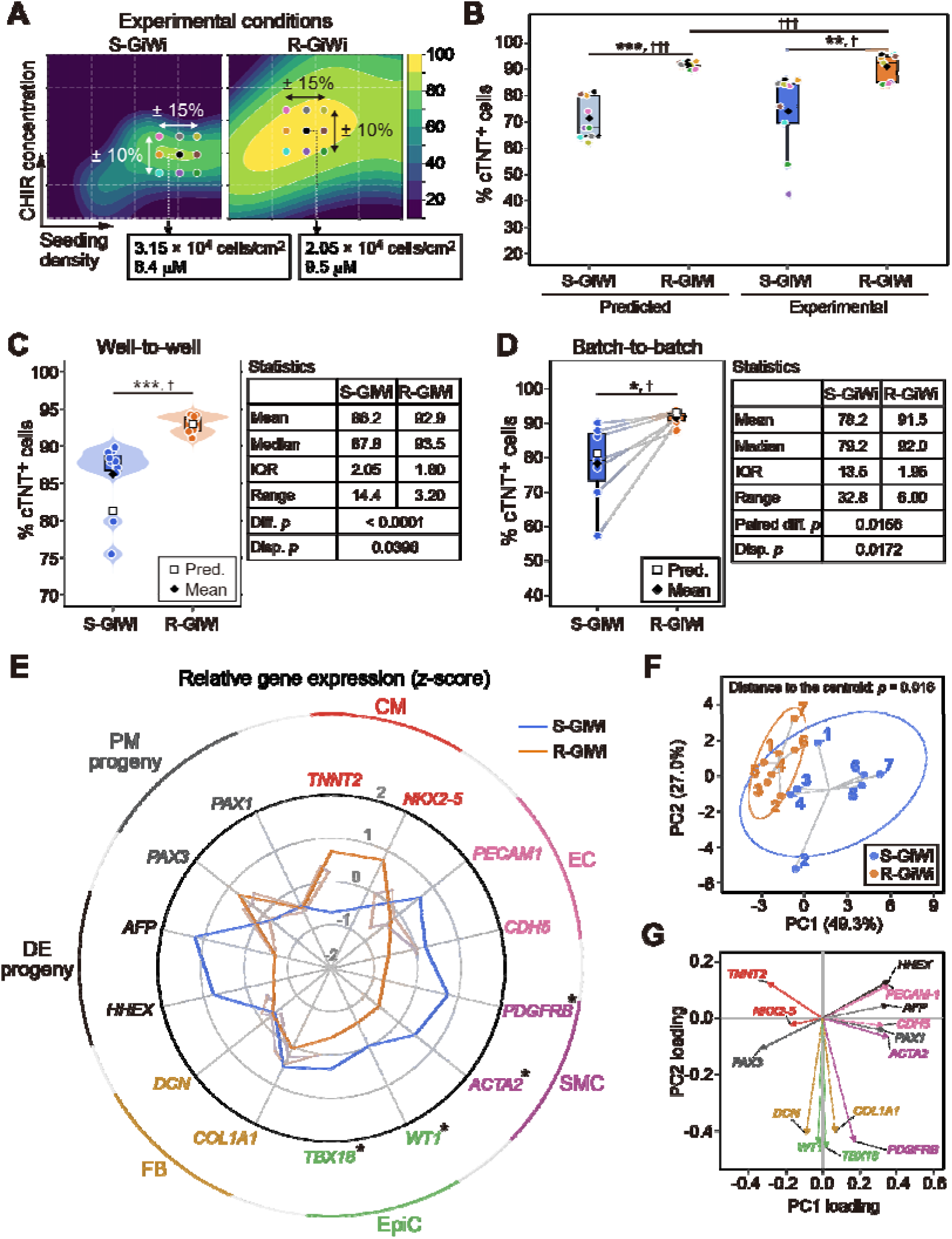
Comparison of process robustness between S-GiWi and R-GiWi. **A:** Experimental conditions used to assess robustness. The predicted optimal condition for each protocol was selected at approximately the center of its high-efficiency window and is indicated by black circles. Eight additional conditions (colored circles) were selected by perturbing the CHIR concentration by ±10% and cell-seeding density by ±15% relative to the corresponding values in the predicted optimal conditions. **B:** Comparison of robustness to parameter perturbations across the nine conditions shown in (A). Predicted and experimentally measured percentages of cTNT-positive cells are shown. Point colors correspond to the conditions shown in (A). Differences in central tendency and dispersion were assessed between S-GiWi and R-GiWi for predicted and experimental values, separately. Predicted and experimental values were also compared within each protocol, using the statistical tests described at the end of this legend. **C:** Comparison of well-to-well performance based on the percentage of cTNT-positive cells under the predicted optimal conditions (n = 11 wells). Open squares indicate predicted values (Pred.), and filled diamonds indicate mean experimental values. Range, minimum to maximum. “Diff. *p*” and “Disp. *p*” denote *p*-values for differences in central tendency and dispersion, respectively. **D:** Comparison of batch-to-batch performance based on the percentage of cTNT-positive cells under the predicted optimal conditions (n = 7 independent batches). Gray lines connect paired samples from the same batch. Detailed data on the percentages of cTNT-positive cells are provided in Table S2. “Paired diff. *p*” denotes the *p*-value for the difference in central tendency between paired samples. **E:** Relative expression of lineage marker genes measured on day 10 of differentiation using qPCR, shown as gene-wise z-scores. Lines and shaded regions indicate the median and IQR, respectively (n = 7 independent batches). Genes are grouped according to seven lineage categories: CM, EC, SMC, EpiC, FB, DE progeny, and PM progeny. Detailed expression data are provided in Fig. S5. **F:** PCA of the gene-wise z-scored expression profiles of the 14 genes representing the seven lineage categories shown in (E). Ellipses indicate the distributions of samples obtained using each protocol. Labels indicate batch numbers, with identical numbers denoting paired samples from the same batch. Distances to the group centroids were compared using the statistical test described at the end of this legend. **G:** PCA loading plot. Differences in central tendency were assessed using the Mann–Whitney U test for unpaired comparisons in (B) and (C) and the Wilcoxon signed-rank test for paired comparisons in (B) and (D)–(F). Differences in dispersion were assessed using Levene’s test in (B)–(D). \**p* < 0.05, \*\**p* < 0.01, and \*\*\**p* < 0.001 for differences in central tendency; ^†^*p* < 0.05 and ^†††^*p* < 0.001 for differences in dispersion. Only statistically significant differences are indicated.

Next, we assessed well-to-well variability across 11 wells within a representative batch (Fig. 4C) and batch-to-batch variability across seven independent batches (Fig. 4D). R-GiWi showed significantly higher differentiation efficiency and lower variability than those observed with S-GiWi in both the well-to-well and batch-to-batch comparisons. Notably, R-GiWi consistently achieved higher differentiation efficiency than that obtained with S-GiWi across all seven batches (Table S2). Even in a batch in which S-GiWi yielded an efficiency below 60%, R-GiWi achieved an efficiency above 90%. Overall, R-GiWi achieved efficiencies above 90% in six of the seven batches, with even the lowest efficiency reaching 87.7%. Collectively, these results demonstrate that R-GiWi achieves higher differentiation efficiency while minimizing both well-to-well and batch-to-batch variability, thereby enabling highly robust CM differentiation.

Furthermore, we examined potential lineage deviations in cells generated across the seven batches by expression analysis of markers associated with cardiac mesoderm, DE, and PM progenies (Figs. 4E and S5). Expression of the CM markers (*TNNT2* and *NKX2-5*) was higher and relatively less variable with R-GiWi than with S-GiWi, consistent with the differentiation efficiency data (Fig. 4D). Notably, endothelial cell (EC) markers (*PECAM1* and *CDH5*) showed decreasing trends, and smooth muscle cell (SMC) markers (*PDGFRB* and *ACTA2*) and epicardial cell (EpiC) markers (*WT1* and *TBX18*) were significantly reduced. Although markers associated with fibroblasts (FBs; *COL1A1* and *DCN*), which may also arise from cardiac mesoderm, showed no clear trends, the overall expression patterns of cardiac mesoderm progeny markers suggest that R-GiWi reduces deviations toward non-CM cardiac lineages. This may also have contributed to both the higher differentiation efficiency and reduced variability observed with R-GiWi. The reduced expression of DE progeny markers (*AFP* and *HHEX*) was consistent with the TF expression analysis during early differentiation (Fig. 2B). Among the PM progeny markers, *PAX1*, a marker of the sclerotome, showed a decreasing trend, whereas *PAX3*, a marker of the dermomyotome, showed an increasing trend. Despite the increasing trend in *PAX3* expression in R-GiWi, the differentiation efficiencies exceeding 90% in most batches, together with the reduced expression of *MEOX1* (Fig. 2B) and *PAX1*, suggest that the overall presence of PM-derived progeny in R-GiWi was likely limited. To comprehensively evaluate these multi-lineage profiles, we performed principal component analysis (PCA) of the expression profiles of these 14 genes representing seven lineages (Fig. 4F and G). R-GiWi showed significantly smaller distances to the centroid, indicating reduced variability in lineage deviation. Overall, these results further support the process robustness of R-GiWi.

### 3.5. CMs generated via an early WNT inhibition–based process exhibit comparable characteristics

Finally, we sought to characterize the molecular and functional properties of CMs from R-GiWi relative to those from S-GiWi. For this purpose, cells from three representative batches of the seven evaluated in the batch-to-batch variability analysis (Fig. 4D) were subjected to metabolic selection to minimize non-CMs and their influence, after which the purified CMs were cultured for an additional 2 weeks to allow further maturation before analyses on day 30 (Fig. 5A and Table S2).

**Fig. 5.**
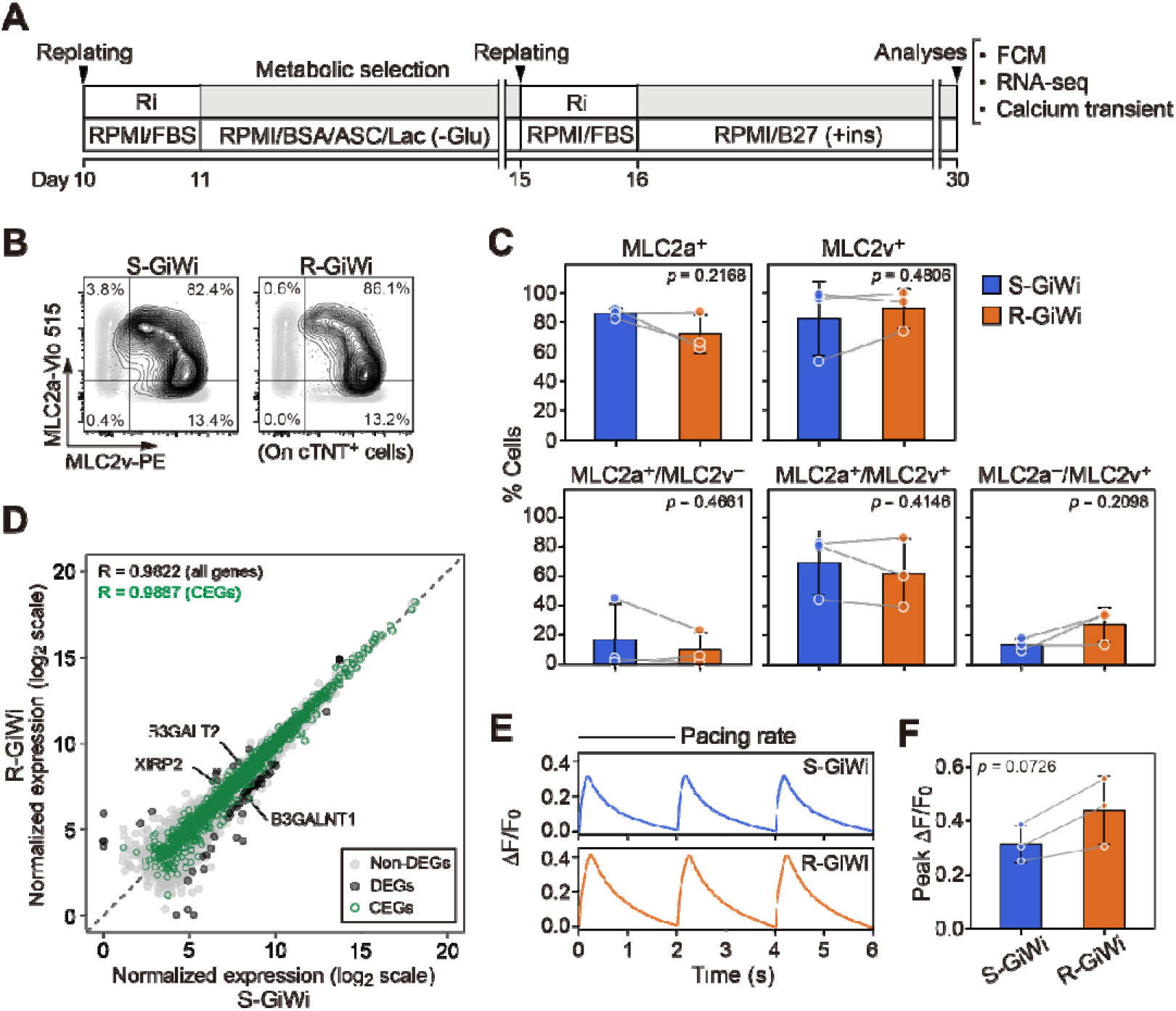
Characterization of CMs derived from the S-GiWi and R-GiWi protocols. **A:** Culture schematic. BSA, bovine serum albumin; ASC, ascorbic acid; Lac, lactate, Glu, glucose. Cells from both protocols were replated on day 10, subjected to metabolic selection for 4 days, and then maintained until day 30, when they were subjected to subsequent analyses. The specific batch allocation of samples used for the analyses is detailed in Table S2. **B:** Representative flow cytometry plots showing the expression of MLC2a and MLC2v in cTNT-positive cells. Light gray contours indicate FMO controls generated from pooled samples. **C:** Comparison of the percentages of the indicated populations. **D:** A scatter plot comparing global gene-expression profiles obtained using bulk RNA-seq. Differential expression was assessed using paired DESeq2 analysis. Differentially expressed genes (DEGs) are highlighted in dark gray, whereas CEGs are outlined in green. The three CEGs identified as differentially expressed are labeled in the plot. Summary statistics of the RNA-seq analysis are provided in Table S3. **E:** Representative calcium transient waveforms recorded under electrical pacing. Representative videos of CMs derived from the S-GiWi and R-GiWi protocols are provided as Movies S1 and S2, respectively. **F:** Comparison of the peak ΔF/F_0_ values. All three analyses were performed using three independent batches (n = 3). Data are presented as mean ± SD in (C) and (F), and gray lines connect paired samples from the same batch. Paired comparisons in (C) and (F) were performed using paired *t*-test. The *p*-values are presented in the graphs.

First, we analyzed CM subtypes using flow cytometry based on the expression of MLC2a and MLC2v, which are markers associated with fetal/atrial and ventricular CM phenotypes, respectively. This analysis was performed to determine whether early WNT inhibition affected CM subtype composition, as cardiac progenitor lineages associated with different chamber identities are specified during the early stages of differentiation [73,74]. As shown in Fig. 5B and C, no significant differences were observed between S-GiWi and R-GiWi CMs in any of the five populations classified based on these marker expression patterns. Although the vast majority of cells expressed MLC2a due to phenotypic immaturity, the mean proportion of MLC2v ventricular CMs exceeded 80% in both groups, consistent with previous reports that the GiWi protocol predominantly generates ventricular CMs [75]. This indicates that early WNT inhibition did not substantially alter this ventricular predominance.

Next, transcriptome-wide comparison revealed a strong correlation between S-GiWi and R-GiWi CMs across all genes (R = 0.9822) (Fig. 5D). One S-GiWi sample subjected to this analysis had a CM purity of 93.9%, which was relatively low compared with the purities of other samples, all of which exceeded 97% (Table S2). As the transcriptomic comparison across all genes could potentially be influenced by residual non-CMs, particularly in the S-GiWi sample with lower purity, we focused on CEGs defined using gene sets from the Human Protein Atlas database [53,54] (Fig. S6); these CEGs are highlighted in green in Fig. 5D. Among these genes, only three (labeled directly in Fig. 5D) were identified as differentially expressed, and their overall expression profiles were also highly correlated between S-GiWi and R-GiWi CMs (R = 0.9887). These results suggest that early WNT inhibition had minimal effects on the overall cardiac gene expression profile of the resulting CMs.

Finally, we assessed the functional properties of the CMs by measuring calcium transients under electrical pacing. CMs from both protocols exhibited regular calcium transients synchronized with the pacing stimuli (Fig. 5E and Movies S1 and S2). The peak calcium transient amplitude tended to be higher in R-GiWi CMs than in S-GiWi CMs, although the difference did not reach statistical significance (Fig. 5F), indicating that early WNT inhibition did not impair calcium-handling function. Taken together, S-GiWi and R-GiWi CMs showed no substantial overall differences in their molecular or functional properties within the scope of the analyses performed, suggesting that they were largely comparable.

## 4. Discussion

In this study, we demonstrated that early WNT inhibition expanded the effective CHIR concentration window toward higher concentrations for CM differentiation across distinct hiPSC lines. Extending this analysis to a two-dimensional process window incorporating cell density further revealed a substantially larger process window for R-GiWi than for S-GiWi. We also characterized temporal changes in lineage-specific TF expression and found that early WNT inhibition suppressed key TFs associated with competing PM and DE fates under conditions of high initial WNT activation. PM suppression appeared to underlie process window expansion, whereas DE suppression may have contributed to more efficient CM differentiation. R-GiWi exhibited greater process robustness to parameter perturbations and variations across wells and batches, along with enhanced differentiation efficiency. Importantly, these process improvements were achieved without compromising the properties of the resulting CMs, yielding CMs with molecular and functional characteristics comparable to those obtained with the standard protocol.

Previous studies have reported conflicting effects of early WNT inhibition on CM differentiation in protocols involving 1 day of CHIR treatment, with some studies showing reduced differentiation efficiency [15,21] and others demonstrating enhanced efficiency [17,25]. In this study, we reconciled these contradictory findings by examining early WNT inhibition across a range of CHIR concentrations and demonstrating that its effects are concentration-dependent. Early WNT inhibition reduced CM differentiation efficiency at lower CHIR concentrations but enhanced it at higher concentrations compared with that achieved with standard delayed WNT inhibition, with this relationship particularly evident in the 253G1 cell line and a similar trend observed in the 610B1 cell line (Fig. 1B, C, F, and G). Primitive streak progression has been shown to depend on the magnitude of CHIR-mediated WNT activation [37]. Accordingly, at lower CHIR concentrations, cells may not have progressed sufficiently along the primitive streak trajectory by day 1 for early WNT inhibition to support subsequent LPM/cardiac specification. Therefore, early WNT inhibition may impair differentiation under these conditions. In S-GiWi, endogenous WNT activity after CHIR withdrawal may allow further progression toward a cardiac-permissive state, resulting in higher differentiation efficiency. In contrast, at higher CHIR concentrations, cells may already be sufficiently advanced by day 1 for WNT inhibition to favor subsequent LPM/cardiac specification, making early WNT inhibition advantageous by preventing excessive posterior progression and thereby increasing differentiation efficiency. This concentration-dependent behavior may account for the conflicting results reported in previous studies. Thus, our findings underscore the difficulty of appropriately evaluating process modifications at a single CHIR concentration. They also raise a broader practical consideration for developing differentiation processes for CMs as well as other primitive streak–derived lineages: modifications involving the timing and/or duration of differentiation factors, medium composition, or compound addition, particularly during early differentiation, should be assessed across a range of CHIR concentrations to avoid potentially misleading conclusions.

Here, we demonstrated the expansion of the CHIR concentration window (Fig. 1) as well as a two-dimensional process window incorporating cell density as an additional critical parameter (Fig. 3), with the window predominantly extending toward higher CHIR concentrations and lower cell densities. The expansion of the CHIR window alone can be explained by early WNT inhibition suppressing stronger WNT-driven posterior mesodermal progression at higher CHIR concentrations, thereby extending the initial WNT activation range that supports LPM/cardiac fate rather than PM specification. To understand how cell density integrates into this relationship, we consider the mechanism demonstrated by Kempf *et al*. [37], who showed that CHIR concentration and cell density jointly regulate primitive streak development and subsequent lineage patterning from DE through LPM to PM fates. Specifically, higher cell densities require greater CHIR exposure to achieve comparable mesodermal patterning because density-dependent paracrine signals restrain posterior shifts in lineage commitment. Our results were broadly consistent with this relationship, showing that the CHIR concentration for efficient CM differentiation generally increases with cell density (Fig. 3C–E). Accordingly, at lower cell densities, the process window extended at lower CHIR concentrations than at higher cell densities. Beyond this baseline trend, the expansion of the effective cell-density range toward lower cell densities with early WNT inhibition was particularly pronounced, a finding consistent with a previous report demonstrating that early WNT inhibition substantially improves CM differentiation even at approximately 1% confluency [25]. This markedly enhanced efficiency at low densities can be interpreted within the same developmental framework: although weaker paracrine signaling at lower cell densities is likely to accelerate posterior mesodermal patterning, early WNT inhibition may effectively limit this trajectory through the same mechanism proposed for the expansion of the CHIR window. Thus, preferential specification of LPM/cardiac over PM fates by early WNT inhibition may provide a unifying mechanism for the substantial two-dimensional expansion.

To understand at the molecular level how early WNT inhibition favors LPM/cardiac fate by suppressing PM specification, we focused on transcriptional regulators during early differentiation. Among *MSGN1* and *TBX6*, two key TFs within an interconnected regulatory network that drives PM specification [63–67], *MSGN1* was strongly suppressed by early WNT inhibition, whereas *TBX6* expression was largely maintained (Fig. 2B). Although Loh *et al*. [40] reported downregulation of both *MSGN1* and *TBX6* expression following WNT inhibition, their cytokine-containing differentiation system may have created a signaling environment distinct from that of the relatively simple GiWi system used here. As loss of either *TBX6* or *MSGN1* severely disrupts PM formation [65], the rapid downregulation of *MSGN1* expression by early WNT inhibition may have impaired PM specification, representing a key mechanism underlying PM suppression in our system. *MSGN1* has been reported as a direct transcriptional target of WNT/β-catenin signaling through cis-regulatory elements containing TCF/LEF-binding sites [66,76], providing a plausible explanation for the rapid decline in *MSGN1* expression following WNT inhibition. Meanwhile, the limited CHIR window observed with the 2-day CHIR followed immediately by 2-day WNT inhibition protocol (Fig. S1) suggests that the duration of *MSGN1* expression may critically influence PM specification and that sufficiently early suppression of *MSGN1* may also be key to robust PM suppression. Moreover, given that early WNT inhibition did not directly promote the expression of tested LPM or cardiac mesoderm TFs and instead modestly delayed *FOXF1* and *NKX2-5* expression (Fig. 2B), possibly because early WNT inhibition also suppressed their upstream regulators, improved CM differentiation is unlikely to reflect direct activation of the LPM/cardiac mesoderm program. Rather, the greater sensitivity of *MSGN1* to WNT inhibition may have preferentially suppressed the PM program, thereby shifting the relative lineage balance toward the LPM/cardiac fate. These findings provide a molecular basis for how early WNT inhibition substantially expands the cardiac process window into higher-CHIR/lower-density regions that would otherwise favor PM specification (Fig. 6).

**Fig. 6.**
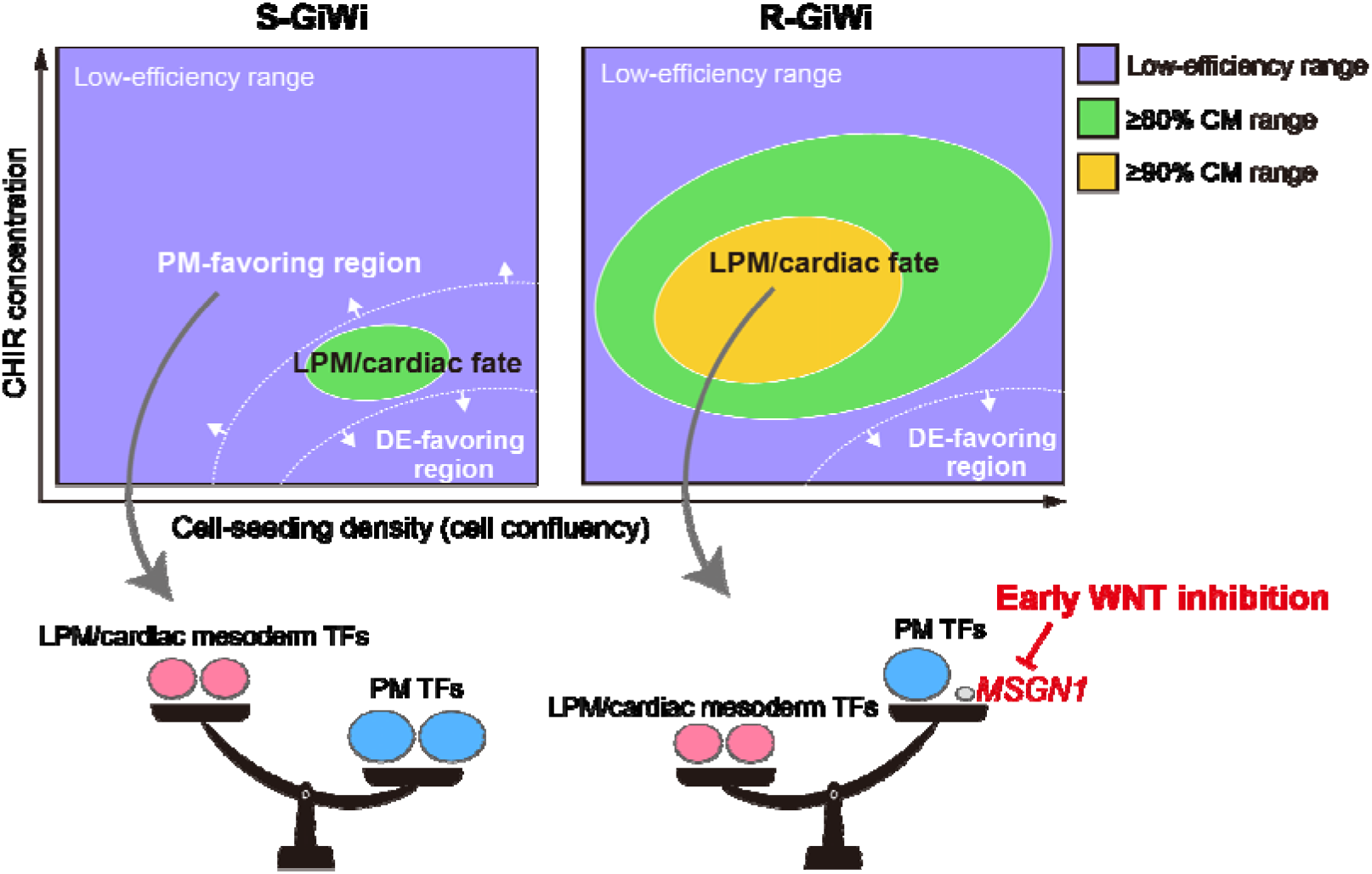
Proposed mechanism underlying the expansion of the process window in cardiac differentiation by R-GiWi.

We assessed the process robustness of R-GiWi from multiple perspectives, including parameter perturbations and well-to-well and batch-to-batch variability, and observed markedly reduced variability in CM differentiation efficiency and consistently higher efficiency than that observed with S-GiWi (Fig. 4). These results support a relationship between process window expansion and the observed improvement in process robustness, reflecting a fundamental principle of process engineering that a broader window can buffer variations in process inputs. From the perspective of cell manufacturing in regenerative medicine, robustness against batch-to-batch variability is particularly important for ensuring consistent production outcomes. In this context, a key observation is that both the high-efficiency range and response peak shifted within the CHIR concentration–cell density parameter space between independent experiments (Fig. 3C and D). Importantly, S-GiWi was still capable of achieving ≥80% efficiency under some conditions in all experiments in which multiple parameter settings were tested (Figs. 1B–C and 3C–D), indicating that the batch-to-batch variability subsequently observed at a fixed condition is unlikely to be explained simply by a loss or reduction in intrinsic differentiation potential of the cells. Rather, it likely reflects batch-dependent changes in cellular responsiveness, with resultant batch-dependent shifts in the critical parameter levels required for appropriate WNT signaling activation. Although we minimized known sources of variation by maintaining a consistent protocol, passage history (e.g., passage number and frequency), and reagent lots (e.g., CHIR and Matrigel) across these two experiments, such shifts still occurred. This suggests that minor technical or biological variations may alter cellular responsiveness in ways that are difficult to fully control in practice. Many studies on stem cell differentiation have predominantly focused on maximizing differentiation efficiency; however, increasing efficiency alone does not necessarily ensure robustness to such batch-dependent shifts. Such shifts may become more likely in large-scale or continuous manufacturing, where operational complexity and changes in material lots introduce additional sources of variability. Thus, our results highlight the importance of process window expansion in directly addressing this issue, as demonstrated here by early WNT inhibition in CM differentiation, with potential relevance to broader stem cell bioprocesses for regenerative medicine.

Another important consideration is what contributes to the higher differentiation efficiency achieved with R-GiWi. Although suppression of PM specification by early WNT inhibition was key to expanding the process window, PM suppression appeared to contribute less directly to the higher efficiency observed with R-GiWi than to that achieved with S-GiWi. This is because gene expression analyses during the early stage of differentiation (Fig. 2B), as well as on day 10 (Fig. 4E), did not provide clear evidence that PM differentiation was more strongly suppressed in R-GiWi than in S-GiWi, because S-GiWi was less prone to PM differentiation under weaker initial WNT activation. Instead, suppression of DE differentiation under stronger initial WNT activation may have contributed more directly to the efficiency improvement, as DE-associated markers were consistently reduced in R-GiWi at both stages (Figs. 2B and 4E). Notably, markers associated with cardiac non-CM lineages, specifically ECs, SMCs, and EpiCs, were reduced in R-GiWi (Fig. 4E), suggesting that suppression of these lineages may also have contributed to the higher differentiation efficiency. These cell types are known to arise from cardiac mesoderm [22,77], which in the GiWi differentiation trajectory corresponds to around day 3 of differentiation, as supported by the peak expression of *MESP1*, one important marker of early cardiac mesoderm [78], observed on day 3 in our experiments (Fig. 2B). The critical window for subsequent specification therefore occurs during days 3–5. In this study, differentiation efficiency progressively increased as the duration of WNT inhibition was extended from days 1–3 to days 1–4 and 1–5 (Fig. 1J), indicating that coverage of the days 3–5 cardiac lineage specification window is advantageous for improving CM differentiation efficiency.

Because S-GiWi also includes WNT inhibition during days 3–5, suppression of cardiac non-CM lineages is unlikely to be explained simply by WNT inhibition during this period. Rather, continuous WNT inhibition beginning at the primitive streak stage and extending beyond the cardiac mesoderm stage may have more effectively suppressed cardiac non-CM lineage specification and thereby favored CM specification. Although early WNT inhibition in previous GiWi-based studies has generally been limited to days 1–3 [17,25], extending inhibition to days 1–5 in the present study therefore represents a further modification of the GiWi strategy, contributing not only to process window expansion but also to higher CM differentiation efficiency. Notably, the optimal CHIR concentration and cell density differed between S-GiWi and R-GiWi (Fig. 4A). The aforementioned studies generally compared WNT inhibition initiated from day 1 versus day 3 under matched CHIR and cell-density conditions [17,25], which may not fully reflect the relative performance of the two protocols. By evaluating both protocols under their respective favorable conditions, the present comparison more accurately reflects their relative differentiation performance and provides a more informative comparison of differentiation efficiency between R-GiWi and S-GiWi.

Line-to-line variability is also an important consideration in CM differentiation [16,19–21]. In the present study, R-GiWi reproducibly achieved approximately 90% differentiation efficiency across batches for 610B1 cells, and it also maintained similar efficiency across a broad CHIR concentration range in 253G1 cells, suggesting that the observed advantages of R-GiWi may extend beyond a single cell line. Nevertheless, validation across a larger number of iPSC lines will be required to establish its broader generalizability. Importantly, we speculate that some of the apparent line-to-line variability observed with conventional protocols may partly arise from insufficient parameter optimization due to the limited window supporting high-efficiency differentiation, as demonstrated here for S-GiWi. In particular, the interdependent relationship between CHIR concentration and cell density, together with batch-dependent shifts in the optimal parameter levels, makes identification of truly favorable conditions difficult using conventional one-factor-at-a-time optimization with a limited number of experiments. Consequently, some cell lines may be evaluated under suboptimal or inherently unstable conditions, potentially exaggerating apparent differences in intrinsic differentiation propensity among lines. By substantially broadening the process window while also improving differentiation efficiency, R-GiWi may facilitate the identification of stable, high-efficiency conditions for individual cell lines and thereby help minimize apparent line-to-line variability in differentiation outcomes.

R-GiWi offers distinct practical advantages for CM manufacturing in regenerative medicine beyond improving process robustness. Quality by Design (QbD) has become an increasingly important framework for manufacturing process development in biopharmaceuticals as well as in cell and tissue products, with the establishment of a design space as one of its central components [32,33,79]. A design space represents the multidimensional combination of input variables and process parameters within which predefined critical quality attributes (CQAs) can be consistently maintained within acceptable ranges. Importantly, independent design spaces can be established for individual unit operations, while a single design space can also span multiple operations or the entire manufacturing process [79]. Establishing a design space could increase operational flexibility, and the associated process understanding could provide a scientific basis for continual process improvement. The process window defined in this study was based solely on the percentage of cTNT-positive cells and may not necessarily capture all CQAs relevant to the differentiation step as a unit operation within the manufacturing process of PSC-derived CMs or cardiac tissues. Nevertheless, the purity of target cells is generally considered one of the fundamental CQAs in cell manufacturing. Therefore, expanding the parameter space enabling high-efficiency CM differentiation with R-GiWi may provide a strong basis for establishing a broader design space within a QbD framework, thereby supporting broader applications of cardiac regenerative medicine.

Recently, a conceptually similar expansion of the effective CHIR concentration window was reported by Yang *et al*. using CDK8 inhibitors [20]. When administered during the 2-day CHIR treatment, these inhibitors broadened the effective range toward higher CHIR concentrations across PSC lines and batches, accompanied by transcriptomic changes consistent with suppression of presomitic mesoderm differentiation under high-CHIR conditions. In contrast, the WNT inhibitor IWR-1 did not reproduce the same degree of window expansion under the conditions tested, suggesting that the effect of CDK8 inhibition cannot be explained solely by attenuation of excessive WNT signaling and may involve additional mechanisms. Although the study by Yang *et al*. provides an important precedent for expanding the effective CHIR concentration window, the mechanism by which CDK8 inhibition suppresses presomitic mesoderm differentiation remains incompletely understood and requires further investigation. Because R-GiWi requires only a change in the timing of WNT inhibition already used in the S-GiWi protocol, it may be more readily adopted by users of GiWi-based cardiac differentiation protocols.

One limitation of this study is that a systematic assessment of batch-to-batch variability in CM quality attributes other than cTNT expression was beyond its scope. Although molecular and functional analyses indicated that CMs generated by R-GiWi were broadly comparable to those generated by S-GiWi (Fig. 5), these analyses were performed on a limited number of representative batches and were therefore insufficient to assess batch-to-batch variability in CM quality. However, the reduced variability in lineage composition achieved with R-GiWi may also be advantageous for minimizing batch-dependent differences in the cellular environment during differentiation, potentially leading to more consistent CM quality. From a more practical perspective, another limitation is that the differentiation system used in this study relied on B27 supplement and Matrigel and was therefore not fully chemically defined, which may limit direct translation to practical manufacturing settings in regenerative medicine. GiWi-based cardiac differentiation has also been demonstrated under chemically defined conditions using chemically defined media such as CDM3 together with recombinant matrices [58]. Thus, future studies should determine whether R-GiWi maintains a substantially expanded process window under fully chemically defined conditions while also reducing variability in CM quality. Despite these limitations, we believe that the substantial process window expansion demonstrated with R-GiWi supports its practical relevance for engineering robust CM differentiation processes.

## 5. Conclusions

We demonstrated that early WNT inhibition can substantially expand the multidimensional process window in GiWi-based CM differentiation while also increasing differentiation efficiency through suppression of alternative cell fates with closely related developmental signaling requirements. Furthermore, assessment of process robustness supported a link between a larger process window and improved robustness, consistent with a fundamental principle of process engineering. Despite the inherently variable nature of stem cell differentiation, systematic understanding and strategic expansion of the process window have been largely overlooked. However, our findings highlight their critical importance for engineering robust differentiation processes. Integrating this perspective into the stem cell field may contribute to establishing more robust and reproducible processes for the generation of CMs and other cell types of interest, with potential benefits ranging from practical cell manufacturing for regenerative medicine to routine stem cell differentiation for biological research.

## CrediT authorship contribution statement

Conceptualization: **H.A.**; Data curation: **H.A.** and **R.K.**; Formal analysis: **H.A.** and **R.K.**; Funding acquisition: **H.A.**; Investigation: **H.A.** and **R.K.**; Methodology: **H.A**., **R.K.**, and **S.N.**; Project administration: **H.A.**; Resources: **H.A.**, **K.S.**, and **H.H.**; Software: **H.A.**; Supervision: **H.A.**, **T.Y.**, **K.S.**, and **H.H.**; Validation: **H.A.** and **R.K.**; Visualization: **H.A.**; Writing – original draft: **H.A.**; Writing – review and editing: **H.A.**, **T.Y.**, and **K.S.** All authors read and approved the final manuscript.

## Declaration of generative AI and AI-assisted technologies in the manuscript preparation process

During the preparation of this work, the authors used ChatGPT (OpenAI) to assist with language editing and improving the clarity and readability of the manuscript. After using this tool, the authors reviewed and edited the content as needed and take full responsibility for the content of the published article.

## Acknowledgements

The authors gratefully acknowledge Mika Nomoto, Yasuomi Tada, Akiko Akama, and Mikako Yamaguchi of the Center for Gene Research, Nagoya University, for their technical support for next-generation sequencing using the NextSeq 550 system. The authors also thank the Division for Medical Research Engineering, Nagoya University Graduate School of Medicine, for providing access to the BD FACSCanto II flow cytometer and FlowJo software. The authors also thank Editage for English language editing of this manuscript.

## Data availability

The RNA-seq data generated in this study have been deposited in the DDBJ Sequence Read Archive under BioProject accession number PRJDB45686. The other datasets and codes generated and/or analyzed during the current study are available from the corresponding author (H.A.) upon reasonable request.

