## Supplementary Information for "Engineering robust cardiomyocyte differentiation from iPSCs through WNT-mediated process window expansion"

* Corresponding author:

**Supplementary methods**

**Cardiac differentiation using 2-day CHIR treatment followed by 2-day WNT inhibition**

iPSCs (610B1) were seeded on Matrigel-coated 48-well plates at a density of 2.2 × 10^4^ cells/cm^2^ in mTeSR Plus medium supplemented with 10 μM Y-27632 and expanded for 3 days with daily medium replacement. The cell-seeding density was identical to that used in the experiments shown in Fig. 1. On day 0 of differentiation, the medium was replaced with RPMI/B27 (−ins) containing 4.5–8.0 μM CHIR. On day 2, the medium was replaced with RPMI/B27 (−ins) containing 5 μM IWP-2. On day 4, the medium was replaced with RPMI/B27 (−ins) without IWP-2. From day 6 onward, the cells were cultured in RPMI/B27 (+ins). On day 10, the cells were harvested using TrypLE Express and analyzed using flow cytometry as described in the Materials and Methods.

**Generation of a cardiac-enriched gene set from the Human Protein Atlas database**

Three gene sets were obtained from different sections of the Human Protein Atlas database (https://www.proteinatlas.org/) to broadly capture genes enriched in cardiac tissue or CMs: (i) 429 genes classified as having elevated expression in heart muscle compared with that in other tissue types, based on tissue-level bulk RNA-seq; (ii) 1,341 genes with a predicted specificity differential score >0.15 for CMs relative to other profiled cell types within heart muscle, based on tissue cell type bulk RNA-seq; and (iii) 1,218 genes classified as having elevated expression in CMs compared with that in other profiled cell types, based on single-cell RNA-seq. These gene sets were combined to generate a nonredundant gene set comprising 2,353 genes (Fig. S6), which was used as a broadly defined cardiac-enriched gene (CEG) set. These genes are indicated by green outlines in Fig. 5D. The gene sets were retrieved from the database on May 28, 2026.

**Supplementary figures**

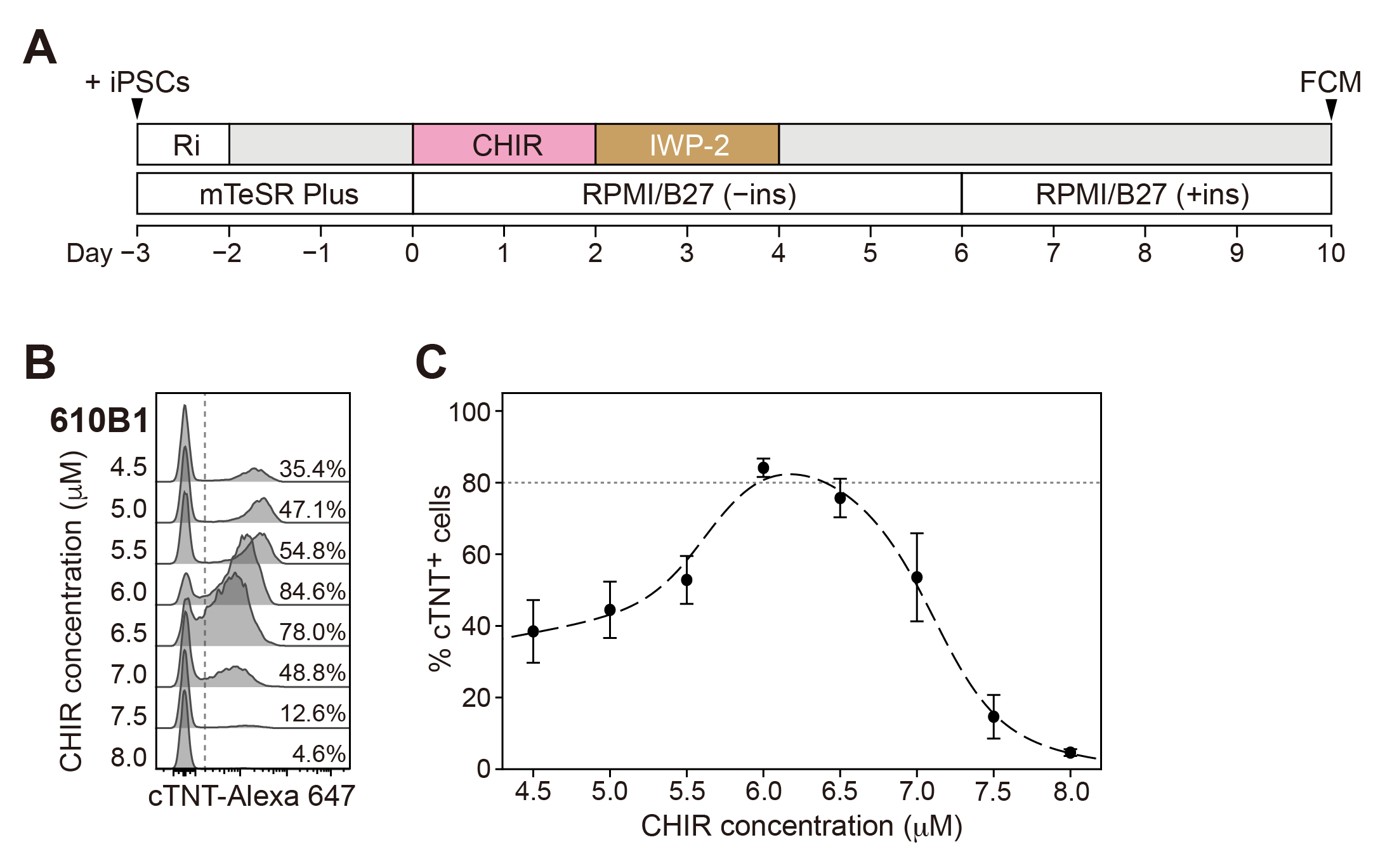
**Fig. S1.** Effect of CHIR concentration on cardiac differentiation efficiency using a GiWi protocol with 2-day CHIR treatment followed by 2-day WNT inhibition. **A:** Schematic of the protocol. **B:** Representative flow cytometry histograms for cTNT expression on day 10 of differentiation in the 610B1 cell line with 4.5−8.0 μM CHIR. **C:** Plots of percentages of cTNT-positive cells on day 10 with 4.5−8.0 μM CHIR. The data were fitted using a GAM; the fitted curve is shown as a dashed line. Data are presented as mean ± SD (n = 3 wells).

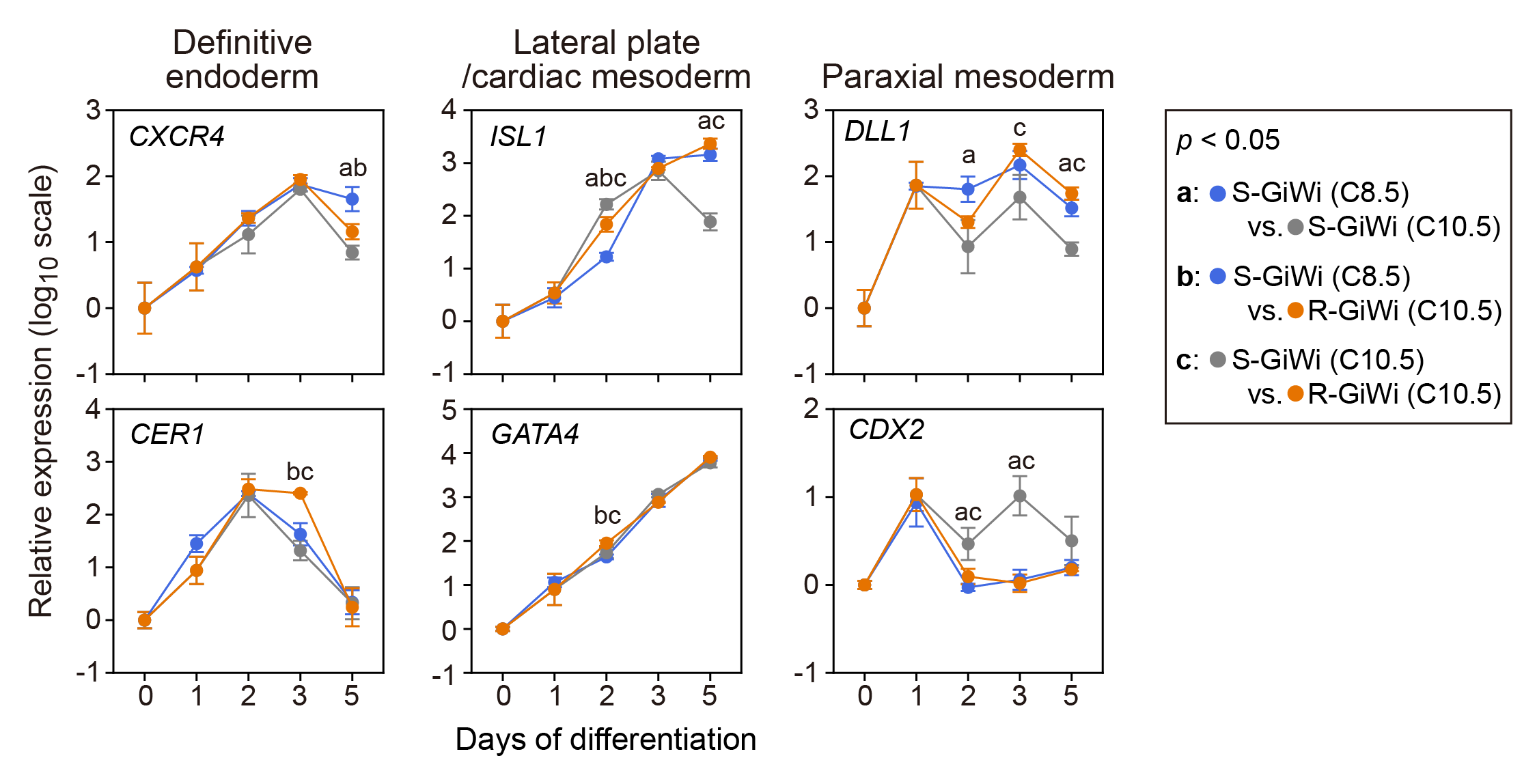

**Fig. S2.** Effects of early WNT inhibition on lineage marker expression during cardiac differentiation. Experimental conditions were the same as those described in Fig. 2. Data are presented as mean ± SD (n = 3 wells). Statistical analyses were performed on days 2, 3, and 5 using one-way ANOVA followed by Tukey’s post hoc test. Letters indicate statistically significant differences (*p* < 0.05), with the corresponding comparisons defined in the figure.

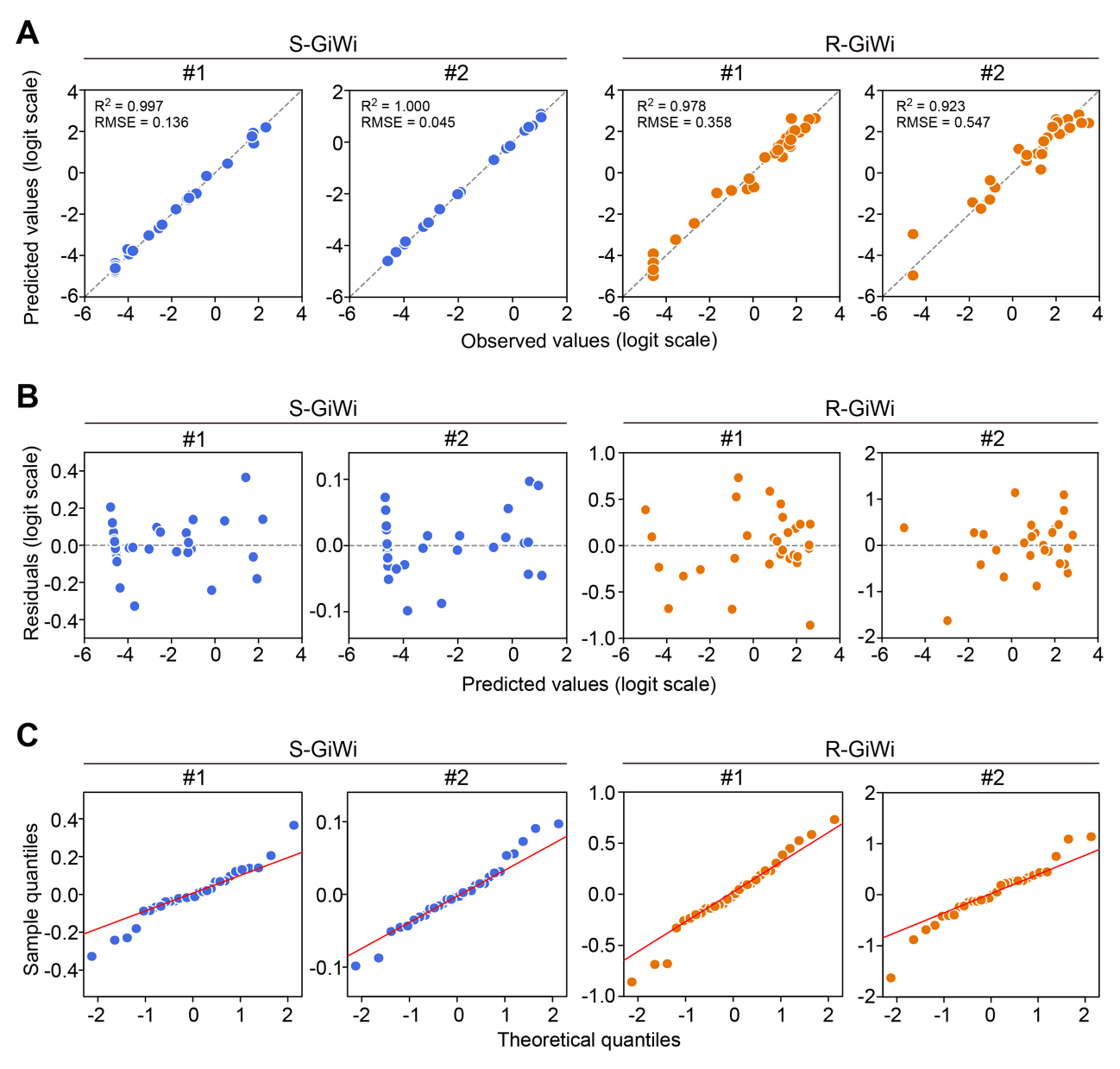

**Fig. S3.** Diagnostic plots for GAMs fitted separately to the response data for each experiment. **A:** Predicted-versus-observed plots for S-GiWi and R-GiWi in experiments #1 and #2. **B:** Residual-versus-predicted plots for the corresponding models. **C:** Normal quantile–quantile plots of the residuals for the corresponding models. Observed and predicted values, as well as residuals, are shown on the logit scale.

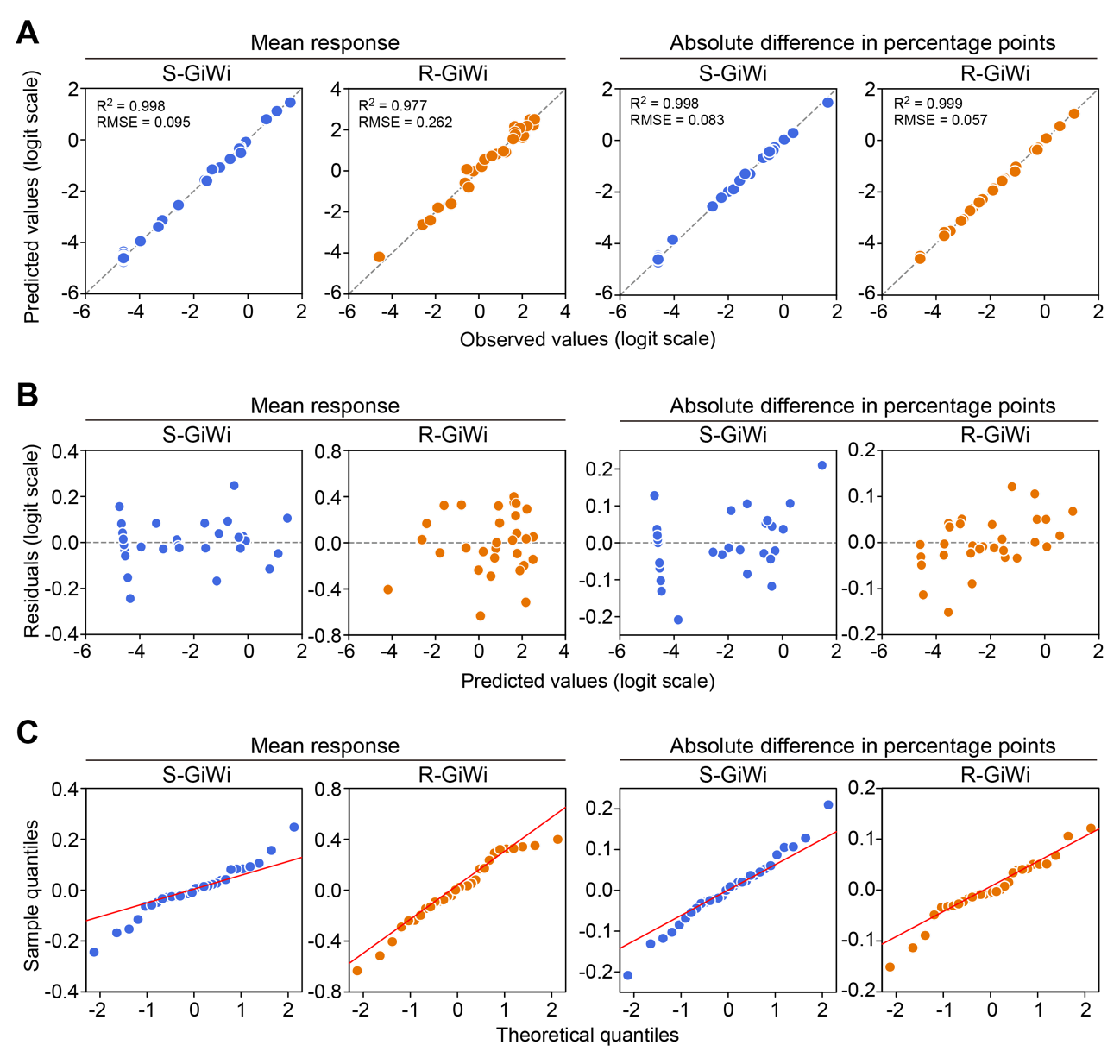

**Fig. S4.** Diagnostic plots for GAMs fitted to the mean response across the two experiments and the between-experiment difference. **A:** Predicted-versus-observed plots for the models of the mean response and between-experiment difference for S-GiWi and R-GiWi. **B:** Residual-versus-predicted plots for the corresponding models. **C:** Normal quantile–quantile plots of the residuals for the corresponding models. Observed and predicted values, as well as residuals, are shown on the logit scale.

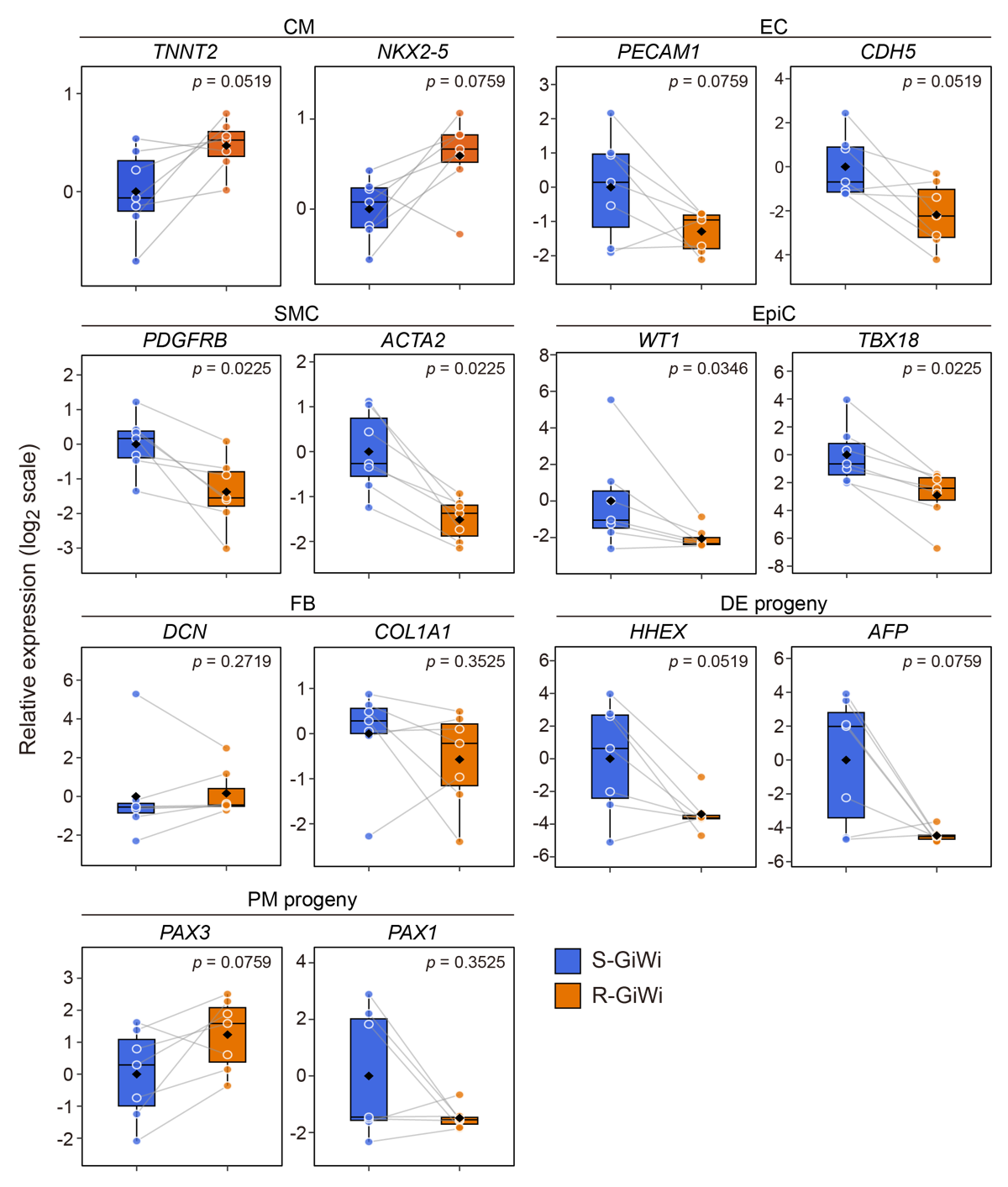

**Fig. S5.** Relative expression of lineage marker genes measured on day 10 using qPCR in S-GiWi and R-GiWi. Detailed relative expression data corresponding to those summarized in Fig. 4E are shown. Gray lines connect paired samples from the same batch, and filled diamonds indicate mean values (n = 7 independent batches). Statistical significance was assessed using the Wilcoxon signed-rank test. The *p*-values are presented in each graph.

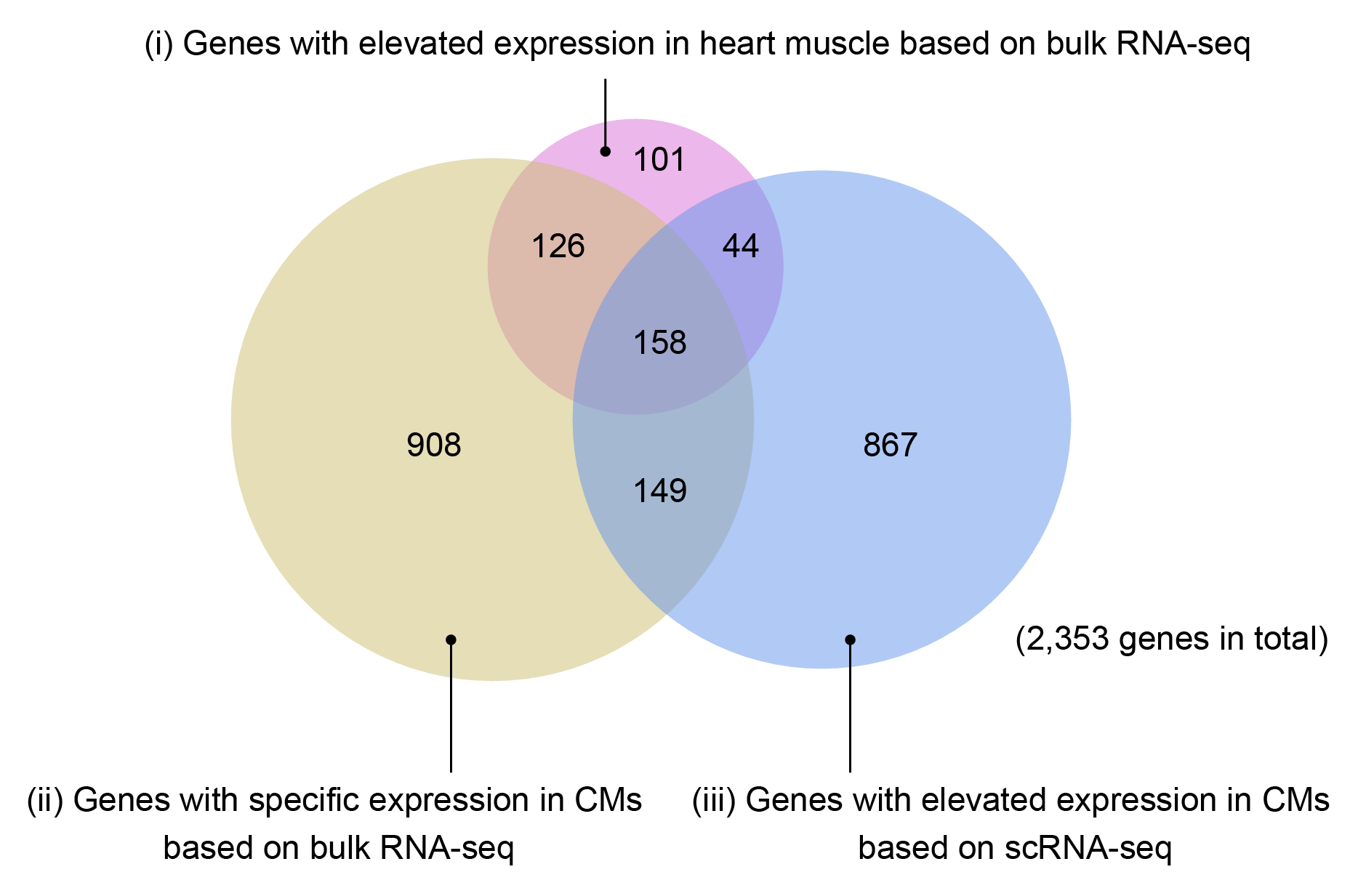

**Fig. S6.** Gene sets used to define CEGs. A nonredundant set of 2,353 genes was generated by combining three gene sets obtained from the Human Protein Atlas database: (i) genes with elevated expression in heart muscle based on tissue-level bulk RNA-seq; (ii) genes with a predicted specificity differential score >0.15 for CMs relative to other profiled cell types within heart muscle, based on tissue cell type bulk RNA-seq; and (iii) genes with elevated expression in CMs compared with that in other profiled cell types, based on single-cell RNA-seq (scRNA-seq), as detailed in the Supplementary Methods.

**Supplementary tables**

**Table S1. List of primers used in this study**

| Gene name |  | 5′–3′ primer sequence | T_m_ (°C) ^a^ | Product size (bp) | Accession # ^b^ |
| --- | --- | --- | --- | --- | --- |
| *ACTA2* | F | CTATGCCTCTGGACGCACAACT | 62.62 | 115 | NM_001141945.3 |
|  | R | CAGATCCAGACGCATGATGGCA | 63.01 |  |  |
| *AFP* | F | GCAGAGGAGATGTGCTGGATTG | 61.32 | 113 | NM_001134.3 |
|  | R | CGTGGTCAGTTTGCAGCATTCTG | 62.97 |  |  |
| *CDH5* | F | GAAGCCTCTGATTGGCACAGTG | 61.77 | 112 | NM_001795.5 |
|  | R | TTTTGTGACTCGGAAGAACTGGC | 61.54 |  |  |
| *CDX2* | F | CGAGCTGAACTTTCCGGCAG | 61.63 | 80 | NM_001265.6 |
|  | R | AGGCTCTCGGATGTTGGTGG | 61.90 |  |  |
| *CER1* | F | CAGGGGGTCATCTTGCCCAT | 61.94 | 140 | NM_005454.3 |
|  | R | GACCCGCATTTCCCAAAGCA | 61.53 |  |  |
| *COL1A1* | F | GATTCCCTGGACCTAAAGGTGC | 60.68 | 107 | NM_000088.4 |
|  | R | AGCCTCTCCATCTTTGCCAGCA | 64.28 |  |  |
| *CXCR4* | F | TCTGTGACCGCTTCTACCCC | 61.26 | 148 | NM_001008540.2 |
|  | R | TTCTGGTGGCCCTTGGAGTG | 62.07 |  |  |
| *DCN* | F | GCTCTCCTACATCCGCATTGCT | 62.75 | 128 | NM_001920.5 |
|  | R | GTCCTTTCAGGCTAGCTGCATC | 61.32 |  |  |
| *DLL1* | F | TGCCTGGATGTGATGAGCAGCA | 64.57 | 110 | NM_005618.4 |
|  | R | ACAGCCTGGATAGCGGATACAC | 61.92 |  |  |
| *EOMES* | F | CTTGTAAGCGAAGGCGGCTG | 61.97 | 119 | NM_001278182.2 |
|  | R | CCCTCCCATGCCTTTTGAGGT | 62.37 |  |  |
| *FOXA2* | F | GGAACACCACTACGCCTTCAAC | 61.69 | 134 | NM_021784.5 |
|  | R | AGTGCATCACCTGTTCGTAGGC | 62.87 |  |  |
| *FOXF1* | F | GCGAGTTCATGTTCGAGGAGG | 61.06 | 115 | NM_001451.3 |
|  | R | AGGTGGTTGAAGCCGAGCC | 62.55 |  |  |
| *GATA4* | F | GCGGTGCTTCCAGCAACTCCA | 65.78 | 139 | NM_001308093.3 |
|  | R | GACATCGCACTGACTGAGAACG | 61.54 |  |  |
| *GATA6* | F | CATCACGGCGGCTTGGATTG | 62.33 | 81 | NM_005257.6 |
|  | R | GTTCACCCTCGGCGTTTCTG | 61.57 |  |  |
| *GSC* | F | GCGATTTGGACTCGGACAGC | 61.70 | 134 | NM_173849.3 |
|  | R | CTCTTTCTCGACCCCCTCCC | 61.33 |  |  |
| *HAND1* | F | TAGCCACCAGCTACATCGCC | 62.03 | 130 | NM_004821.3 |
|  | R | AGCTCCCTTTTCCGCTTGCT | 62.42 |  |  |
| *HHEX* | F | TCTCAATGTTCGCCCTCCCC | 61.91 | 88 | NM_002729.5 |
|  | R | CGCCCTCAATGTCCACTTCC | 61.03 |  |  |
| *ISL1* | F | ACACCTTGCGGACCTGCTAC | 62.45 | 96 | NM_002202.3 |
|  | R | GGATCACACGGGGACTGAGG | 61.96 |  |  |
| *MEOX1* | F | GCCAAAGAGCATGGGGGACT | 62.51 | 80 | NM_004527.4 |
|  | R | ACCTCCCTCTGCAACACCAG | 61.77 |  |  |

| Gene name |  | 5′–3′ primer sequence | T_m_ (°C) ^a^ | Product size (bp) | Accession # ^b^ |
| --- | --- | --- | --- | --- | --- |
| *MESP1* | F | AGCTGCACCCGAGCCGCGC | 71.02 | 132 | NM_018670.4 |
|  | R | ATCCAGGTCTCCAACAGAGCCA | 63.11 |  |  |
| *MESP2* | F | CTGGGCAACACCCCCTTACT | 61.79 | 80 | NM_001039958.2 |
|  | R | AAAGAGACGCGTCGGAGGTT | 61.80 |  |  |
| *MIXL1* | F | ACCCCGTCTCTTCAACCCTC | 61.19 | 125 | NM_001282402.2 |
|  | R | CTAGAGGCAAGGGGGCTGTC | 61.98 |  |  |
| *MSGN1* | F | GCCAGGGAGCCATTCCTTCT | 61.93 | 94 | NM_001105569.3 |
|  | R | GTCAGAGGAGCCCAAGCCAT | 61.92 |  |  |
| *NANOG* | F | TTGGGATTGGGAGGCTTTGCT | 62.31 | 131 | NM_024865.4 |
|  | R | TTAGCACAACCAACAAATTAGGGGA | 60.93 |  |  |
| *NKX2-5* | F | CCAAGTGTGCGTCTGCCTTTC | 62.32 | 89 | NM_004387.4 |
|  | R | TTCGGCTCTAGGGTCCTTGG | 60.98 |  |  |
| *PAX1* | F | CCGCAGTGAATGGGCTAGAGAA | 62.37 | 133 | NM_006192.5 |
|  | R | TACACGCCGTGCTGGTTGGAG | 65.21 |  |  |
| *PAX3* | F | GGCTTTCAACCATCTCATTCCCG | 62.04 | 146 | NM_181457.4 |
|  | R | GTTGAGGTCTGTGAACGGTGCT | 63.00 |  |  |
| *PDGFRB* | F | TGCAGACATCGAGTCCTCCAAC | 62.30 | 109 | NM_002609.4 |
|  | R | GCTTAGCACTGGAGACTCGTTG | 61.24 |  |  |
| *PECAM1* | F | AAGTGGAGTCCAGCCGCATATC | 62.44 | 133 | NM_000442.5 |
|  | R | ATGGAGCAGGACAGGTTCAGTC | 62.00 |  |  |
| *POU5F1* | F | GCCCGAAAGAGAAAGCGAACC | 62.12 | 98 | NM_002701.6 |
|  | R | CTGATCTGCTGCAGTGTGGG | 61.03 |  |  |
| *RPL37A* | F | GTGGTTCCTGCATGAAGACAGTG | 61.91 | 84 | NM_000998.5 |
|  | R | TTCTGATGGCGGACTTTACCG | 60.40 |  |  |
| *SOX17* | F | GGTGGACCGCACGGAATTTG | 62.20 | 83 | NM_022454.4 |
|  | R | CGGAGTCATGCCCCTGGTAG | 62.03 |  |  |
| *TBX18* | F | CACAACCGTCACTGCCTATCAG | 61.25 | 151 | NM_001080508.3 |
|  | R | CCGTAGTGATGGTCGCCAGAAT | 62.42 |  |  |
| *TBX6* | F | CTTTGCCCCGCACTTTCTCC | 61.87 | 108 | NM_004608.4 |
|  | R | GGCCCAGCAGTGGTTCAGTA | 62.12 |  |  |
| *TBXT* | F | GGAGGATGTTTCCGGTGCTG | 61.03 | 180 | NM_003181.4 |
|  | R | AGTCGGGGTGGATGTAGACG | 61.04 |  |  |
| *TNNT2* | F | TTCACCAAAGATCTGCTCCTCGCT | 64.18 | 166 | NM_000364.4 |
|  | R | TTATTACTGGTGTGGAGTGGGTGTGG | 64.34 |  |  |
| *WT1* | F | CGAGAGCGATAACCACACAACG | 61.85 | 138 | NM_000378.6 |
|  | R | GTCTCAGATGCCGACCGTACAA | 62.34 |  |  |

^a^ Melting temperature (T_m_) values were obtained using the NCBI Primer-BLAST tool.

^b^ Accession numbers are provided for the representative isoform of each gene when multiple isoforms are present.

**Table S2. Summary of cTNT-positive cell percentages and experimental allocation**

| Batch # | % cTNT^+^ cells ^a^ | | | |  | Experimental allocation ^c^ | | |
| --- | --- | --- | --- | --- | --- | --- | --- | --- |
|  | Day 10 | | Day 30 ^b^ | |  | Subtype analysis | RNA-seq | Calcium transient |
|  | S-GiWi | R-GiWi | S-GiWi | R-GiWi |  |  |  |  |
| 1 | 86.2 | 92.9 | — | — |  | — | — | — |
| 2 | 57.2 | 90.3 | — | — |  | — | — | — |
| 3 | 90.0 | 92.6 | — | — |  | — | — | — |
| 4 | 87.7 | 93.7 | 93.9 | 99.0 |  | ✓ | ✓ | ✓ |
| 5 | 79.2 | 87.7 | 97.4 | 98.3 |  | ✓ | ✓ | ✓ |
| 6 | 77.0 | 91.2 | 98.1 | 98.2 |  | ✓ | ✓ | ✓ |
| 7 | 69.8 | 92.0 | — | — |  | — | — | — |

^a^ Values are shown as the mean percentage of cTNT-positive cells obtained under the optimal condition in each batch.

^b^ cTNT-positive cells were measured on day 30, with metabolic selection performed from days 11 to 15 of differentiation. A dash indicates not applicable.

^c^ ✓, performed; —, not performed.

**Table S3. Summary statistics for RNA-seq**

| Sample name | Batch # | Raw reads | Post-trimming reads | Uniquely mapped reads | % Uniquely mapped reads |
| --- | --- | --- | --- | --- | --- |
| S-GiWi rep 1 | 4 | 12,877,738 | 12,876,742 | 11,889,733 | 92.3 |
| S-GiWi rep 2 | 5 | 13,299,954 | 13,298,476 | 12,066,757 | 90.7 |
| S-GiWi rep 3 | 6 | 11,759,735 | 11,758,642 | 10,538,148 | 89.6 |
| R-GiWi rep 1 | 4 | 11,151,275 | 11,150,272 | 9,876,979 | 88.6 |
| R-GiWi rep 2 | 5 | 12,192,791 | 12,191,830 | 10,658,462 | 87.4 |
| R-GiWi rep 3 | 6 | 10,585,043 | 10,584,086 | 9,334,943 | 88.2 |
